# Loss of imprinting of the *Igf2-H19* ICR1 enhances placental endocrine capacity via sex-specific alterations in signalling pathways in the mouse

**DOI:** 10.1101/2021.05.14.444241

**Authors:** Bethany R. L. Aykroyd, Simon J. Tunster, Amanda N. Sferruzzi-Perri

## Abstract

Imprinting control region (ICR1) controls the expression of the *Igf2* and *H19* genes in a parent-of-origin specific manner. Appropriate expression of the *Igf2-H19* locus is fundamental for normal fetal development, yet the importance of ICR1 in the placental production of hormones that promote maternal nutrient allocation to the fetus is unknown. To address this, we used a novel mouse model to selectively delete ICR1 in the endocrine junctional zone (Jz) of the mouse placenta (Jz-ΔICR1). The Jz-ΔICR1 mice exhibit increased *Igf2* and decreased *H19* expression specifically in the Jz. This was accompanied by an expansion of Jz endocrine cell types due to enhanced rates of proliferation and increased expression of pregnancy-specific glycoprotein 23 in the placenta of both fetal sexes. However, changes in the endocrine phenotype of the placenta were related sexually-dimorphic alterations to the abundance of IGF2 receptors and downstream signalling pathways (PI3K-AKT and MAPK). There was no effect of Jz-ΔICR1 on the expression of targets of the *H19* embedded miR-675 or on fetal weight. Our results demonstrate that ICR1 controls placental endocrine capacity via sex-dependant changes in signalling.

**Summary statement:** Imprinting at *Igf2-H19* ICR1 regulates endocrine cell formation and function via sexually-dimorphic changes in PI3K-AKT and MAPK signalling in the mouse.

## Introduction

In mammals, a subset of autosomal genes exhibit monoallelic (Ferguson-Smith and Surani, 2001) or preferential expression of one allele (Khatib, 2007; Schulz et al., 2006) in a parent-of-origin dependent manner. The expression of such imprinted genes is regulated by epigenetic mechanisms, such as DNA methylation, chromatin remodelling, and reciprocal expression of long non-coding RNA (Ferguson-Smith, 2011). To date, 260 imprinted genes have been identified in mice and 228 in humans, with the imprinting status of 63 genes conserved in both species (Tucci et al., 2019). Rather than being dispersed throughout the genome, imprinted genes typically co-localise in clusters, or imprinted domains, that are coordinately regulated by an imprinting control region (ICR) or imprinting centre (IC) (Cattanach and Kirk, 1985; Verona et al., 2003).

The first evidence of genomic imprinting came from pioneering nuclear transplantation experiments undertaken in the 1980s. The developmental failure of conceptuses carrying two paternal genomes (androgenetic) or two maternal genomes (parthenogenetic or gynogenetic) established the absolute requirement of both parental genomes for successful feto-placental development (Barton et al., 1984; Mann and Lovell-Badge, 1984; McGrath and Solter, 1984; Surani et al., 1984). Whilst several theories exist that attempt to explain the evolutionary origins of imprinting (reviewed in Edwards et al., 2019), the most prominent is the parental conflict hypothesis (Moore and Haig, 1991). Essential for mammalian viviparous reproduction is the substantial investment of maternal resources, such as the provision of nutrients from mother to fetus throughout gestation. The parental conflict hypothesis theorises that whilst the father (acting through the paternal genome) is primarily interested in achieving maximal offspring growth, the mother (acting through the maternal genome) must balance supporting growth of the offspring with need for nutrients to sustain her own health, and to support future offspring. Consequently, paternally expressed imprinted genes would be expected to promote fetal growth, whilst maternally expressed imprinted genes would be expected to limit growth (Moore and Haig, 1991).

The placenta functions as the interface between mother and fetus during pregnancy, and it is therefore unsurprising that many imprinted genes exert their influence upon fetal growth by regulating growth, development, and function of this vital organ (Tunster et al., 2013). Indeed, placentation and genomic imprinting are thought to have co-evolved ~168 million years ago (reviewed in Kaneko-Ishino and Ishino, 2019). Arguably the primary role of the placenta is to mediate nutrient and oxygen transfer to fetus (Burton and Fowden, 2015; Sferruzzi-Perri and Camm, 2016). However, the placenta is also a major endocrine organ, producing an abundance of hormones and signalling factors that act systemically to adapt maternal physiology, metabolism and behaviour to support fetal growth and sustain pregnancy (Napso et al., 2018). We recently reported that the placental secretome comprises in excess of 300 proteins, including known factors such as steroid hormones, prolactin/placental lactogens and pregnancy specific glycoproteins (PSGs), as well as novel secreted placental proteins (Napso et al., 2020). Secreted placental proteins such as prolactins (PRLs) and steroidogenic hormones act systemically to regulate maternal insulin production, insulin sensitivity/resistance and glucose metabolism (Ahmed-Sorour and Bailey, 1980; Brelje et al., 2004; Huang et al., 2009; Jarrett et al., 1984; Petres and Sferruzzi-Perri, 2021; Sferruzzi-Perri et al., 2020; Wada et al., 2010). Moreover, PSGs can also act locally to promote immune-modulation and angiogenesis to support fetal development (Blois et al., 2012; Snyder et al., 2001). An imbalance in the allocation of nutrients between the mother and fetus has been linked to abnormal in utero development and lifelong health complications for offspring (Camm et al 2018; Fowden et al., 2006; Gluckman et al., 2008; Sferruzzi-Perri et al., 2013a).

In humans, both transport and endocrine functions are performed by syncytiotrophoblast cells of the placenta (Dearden and Ockleford, 1983), whereas in the mouse, these functions are performed by the structurally distinct labyrinth zone (Lz) and junctional zone (Jz), respectively. The Jz primarily comprises three trophoblast lineages; the spongiotrophoblast (SpT), glycogen cells (GC), and trophoblast giant cells (TGC). These Jz cell types derive from a common *Tpbpa-positive* precursor (Lescisin et al., 1988; Simmons et al., 2007) and have the capacity to produce a variety of hormones, including members of the PRL family, steroidogenic hormones and PSGs (Lavoie and King, 2009; McLellan et al., 2005; Simmons et al., 2008). Additionally, GC accumulate stores of glycogen and are considered to be analogous to the extravillous cytotrophoblast cells of the human placenta (Georgiades et al., 2002; Wislocki and Bennett, 1943).

Numerous mouse models exist that establish the vital role for imprinted genes in regulating placental nutrient transport (Angiolini et al., 2006; Coan et al., 2005). Perhaps key amongst these is the paternally expressed *Igf2,* which encodes for Insulin-like Growth Factor 2 (IGF2) and is highly expressed in placental and fetal tissues of humans (Han et al., 1987, 1988) and mice (DeChiara et al., 1991; Sferruzzi-Perri, 2018). Consistent with the parental conflict hypothesis, paternal inheritance of an *Igf2*-null allele restricts feto-placental growth (Baker et al., 1993; DeChiara et al., 1990, 1991). This fetal growth restriction (FGR) can be attributed, at least in part, to a placental defect, with placenta-specific loss of *Igf2* also restricting fetoplacental growth (Constância et al., 2002).

IGF2 is a potent promoter of cellular proliferation and differentiation, acting through the insulin receptor (INSR) or type-1 IGF receptor (IGF1R) to activate the RAS-MAPK-ERK or PI3K-AKT signalling pathways (reviewed in Czech, 1989; Forbes and Westwood, 2008; Jones and Clemmons, 1995; Sferruzzi-Perri et al., 2017; Siddle, 2011). Binding of IGF2 to INSR and activation of the PI3K-AKT signalling pathway also regulates glucose uptake and glycogen synthesis (Cross et al., 1995; Forbes and Westwood, 2008; Huang et al., 2018; Sferruzzi-Perri et al., 2016). IGF2 is thought to be cleared from the circulation via targeted lysosomal degradation following binding to the type-2 IGF receptor (IGF2R) (Czech, 1989; Lau et al., 1994; Morgan et al., 1987), which is encoded by the *Igf2r* gene that in mice is also imprinted, with expression derived from the maternally inherited allele (Barlow et al., 1991).

*Igf2* localises to the ICR1 imprinted domain on chromosome 7 in mice. This domain consists of the paternally expressed *Ins2, Igf2* and *Igf2as,* the maternally expressed long non-coding *H19* RNA and the microRNAs *mir-675* and *mir-483. Igf2* and *H19* share a common enhancer element, with imprinting mediated through paternal methylation of the differentially methylated region (DMR) ICR1 located ~4 kb upstream of *H19* (Ferguson-Smith et al., 1993; Leighton et al., 1995b; Thorvaldsen et al., 1998; Tremblay et al., 1997). ICR1 contains recognition motifs for the zinc-finger DNA-binding protein CTCF, which blocks the interaction with enhancer elements (Bell et al., 1999; Szabo et al., 2000). Binding of CTCF to the hypomethylated maternal ICR1 prevents interaction of the *Igf2* promoter with downstream enhancers, inactivating the maternal *Igf2* allele (Bell and Felsenfeld, 2000; Kanduri et al., 2000; Szabo et al., 2000) and instead promotes the transcription of *H19* (Engel et al., 2006; Schoenherr et al., 2003). In contrast, CTCF is unable to bind to the methylated paternal allele, allowing interaction of the *Igf2* promoter with downstream enhancer elements and enabling its transcription. The absence of CTCF renders the paternal *H19* allele inactive.

Maternal inheritance of a 13 kb deletion spanning *H19* and ICR1 (*H19^Δ13^*) results in fetoplacental overgrowth in mice (Leighton et al., 1995a). However, normalisation of fetal growth in mice inheriting the *H19^Δ13^* allele maternally and an *Igf2* null allele paternally isolates this fetal over-growth to over-expression of *Igf2* rather than loss of function of *H19* (Leighton et al., 1995a). Indeed, maternal inheritance of a ~1.6 kb deletion spanning ICR1 results in reactivation of the maternal *Igf2* allele and overgrowth of neonates relative to control littermates (Thorvaldsen et al., 1998). Much of the subsequent investigation of *Igf2* function in the placenta has focused on its role in regulating nutrient transport function (Angiolini et al., 2011; Coan et al., 2008; Constância et al., 2005; Sibley et al., 2004). However, both global and placenta-specific loss of *Igf2* also restrict Jz size alongside impacting Lz size and function (Coan et al., 2008; Sferruzzi-Perri et al., 2011). Furthermore, *Igf2* is highly expressed by GC (Redline et al., 1993), with constitutive loss of *Igf2* resulting in reduced GC abundance and placental glycogen stores (Lopez et al., 1996), whilst ubiquitous maternal inheritance of the *H19^Δ13^* allele results in an expansion of the GC population and increased placental glycogen content (Esquiliano et al., 2009).

Recent mouse studies demonstrate an emerging role for imprinted genes in regulating placental endocrine capacity (reviewed in John, 2013, 2017). For instance, over-expression of the maternally expressed imprinted genes *Phlda2* and *Ascl2* result in a reduction in Jz size (Tunster et al., 2015, 2016), suggesting that imprinting (paternal silencing) of these genes enhances placental endocrine capacity. Conversely, loss of expression of the paternally expressed *Peg3* also restricts Jz size (Tunster et al., 2018), suggesting that imprinting (maternal silencing) of *Peg3* would act to restrict placental endocrine capacity. We recently reported that Jz-specific loss of *Igf2* restricts placental endocrine capacity in a sexually-dimorphic manner (Aykroyd et al., 2020). We therefore hypothesised that the acquisition of imprinting of the ICR1 domain modulates placental endocrine capacity. Utilising a unique genetic model in which ICR1 is specifically deleted in cells of the placental Jz (Jz-ΔICR1), we sought to investigate the role of ICR1 imprinting in modulating placental endocrine function.

## Results

### Validation of Jz specific *Igf2-H19* imprinted gene dysregulation with Jz-ΔICR1

Homozygous *TpbpaCre* males (Simmons et al., 2007) were mated to heterozygous ICR floxed females (LoxP sites surrounding the ICR; termed ICR1Flox; Srivastava et al., 2000) for a conditional deletion of the *Igf2* and *H19* ICR in the placental Jz (Fig. 1). This generated litters consisting of fetuses with control and Jz-ΔICR1 placentae (Fig. 1B) and using qPCR of isolated Jz obtained on gestational day (D) 16, we verified that Jz expression of *Igf2* was increased by 30% in males and 25% in females, whilst *H19* decreased by 36% in males and 39% in females (Fig. 2A). In contrast, expression of *Igf2* and *H19* was unaltered in the Lz of Jz-ΔICR1 placentae. To ensure that Jz-ΔICR1 did not result in ectopic expression of *Igf2* or *H19,* we assessed their spatial expression by *in situ* hybridisation. Consistent with previous work (Aykroyd et al., 2020; Coan et al., 2006; Redline et al., 1993) we observed high levels of *Igf2* expression in the Lz and GC, with lower levels of expression in SpT and TGC of both control and Jz-ΔICR1 placentae on D16 (Fig. 2B). In control and Jz-ΔICR1 placentae, *H19* was also widely expressed in the Lz, although expression was restricted to the GC in the Jz (Fig. 2C). Negative controls for the in situ hybridisations are shown in Fig. S1A,B.

**Fig. 1.**
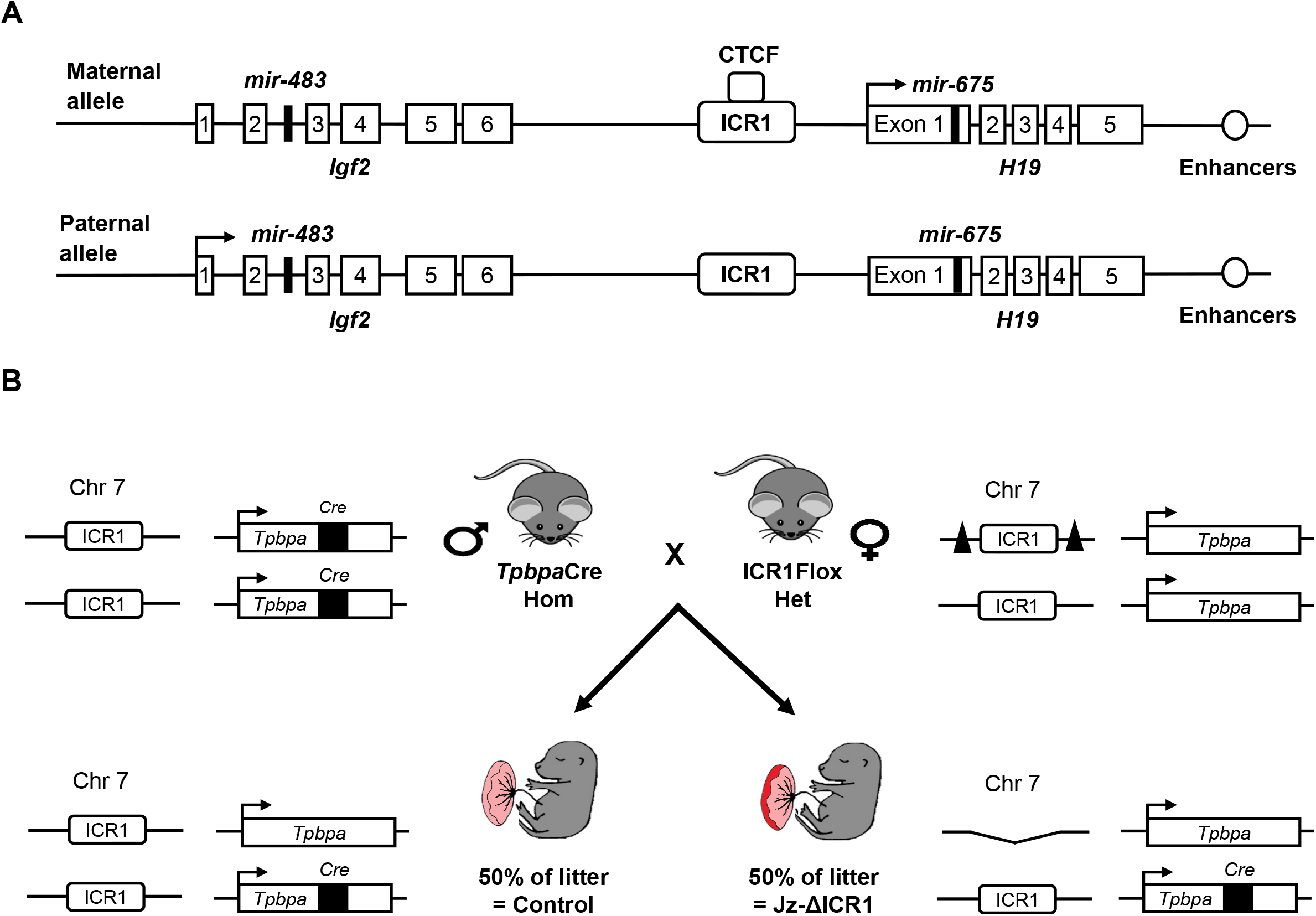
Jz-ΔICR1 mouse genetic manipulation model. (A) Schematic diagram of the *Igf2/H19* gene locus. Imprinting control region (ICR), CCCTC binding factor (CTCF). (B) Breeding strategy to produce litters with control and Jz-ΔICR1 conceptuses. *Cre*-recombinase (*Cre*), chromosome (Chr), homozygous (hom), heterozygous (het), triangles represent lox-P sites and arrows represent transcriptional direction.

**Fig. 2.**
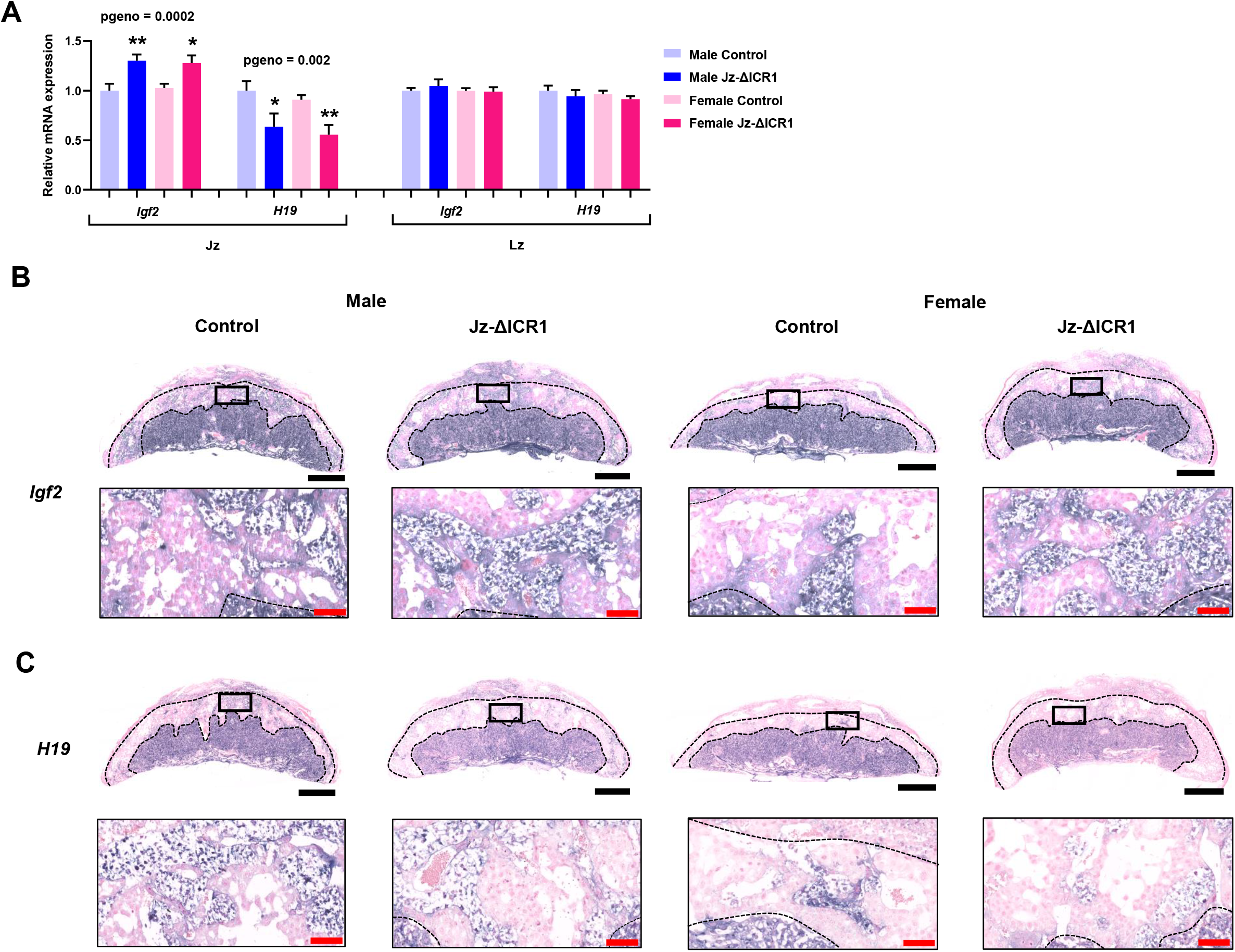
*Igf2* expression is increased and *H19* expression is decreased in the Jz of mouse placentas with Jz-ΔICR1. (A) Expression of *Igf2* and *H19* in isolated Jz and Lz samples on D16 relative to the geometric mean of housekeeping genes (*Hprt* and *Ywhaz* for Jz, *Hprt* and *Polr2a* for Lz) using qPCR (n = 9-10 per genotype/sex in Jz and Lz, across 11 litters). Values presented as mean + SEM with significance assessed by two-way ANOVA and pairwise T-test (pgenotype < 0.05 = *, < 0.01 = **). *In situ* hybridization of (B) *Igf2* and (C) *H19* in males and females. Black boxes represent the area magnified in the image below. Black bar represents 1 mm. Red bar represents 100 μm.

### Jz-ΔICR1 affects placental structure, through enhanced proliferation of endocrine cells, but does not affect not fetal growth

There was no difference in fetal weight (Fig. 3A) or placental weight (Fig. 3B) in response to Jz-ΔICR1. Regardless of genotype, placentae of males were heavier than females (Fig. 3B). Jz-ΔICR1 increased Jz volume by 20% in males and 43% in females. There was an overall effect of Jz-ΔICR1 to decrease Lz and Db volume, with a significant effect of lower Db volume for male conceptuses only (Fig. 3C). Jz volume was lower in placentae of females compared to males, an effect significant in control but not Jz-ΔICR1 conceptuses (Fig. 3C).

**Fig. 3.**
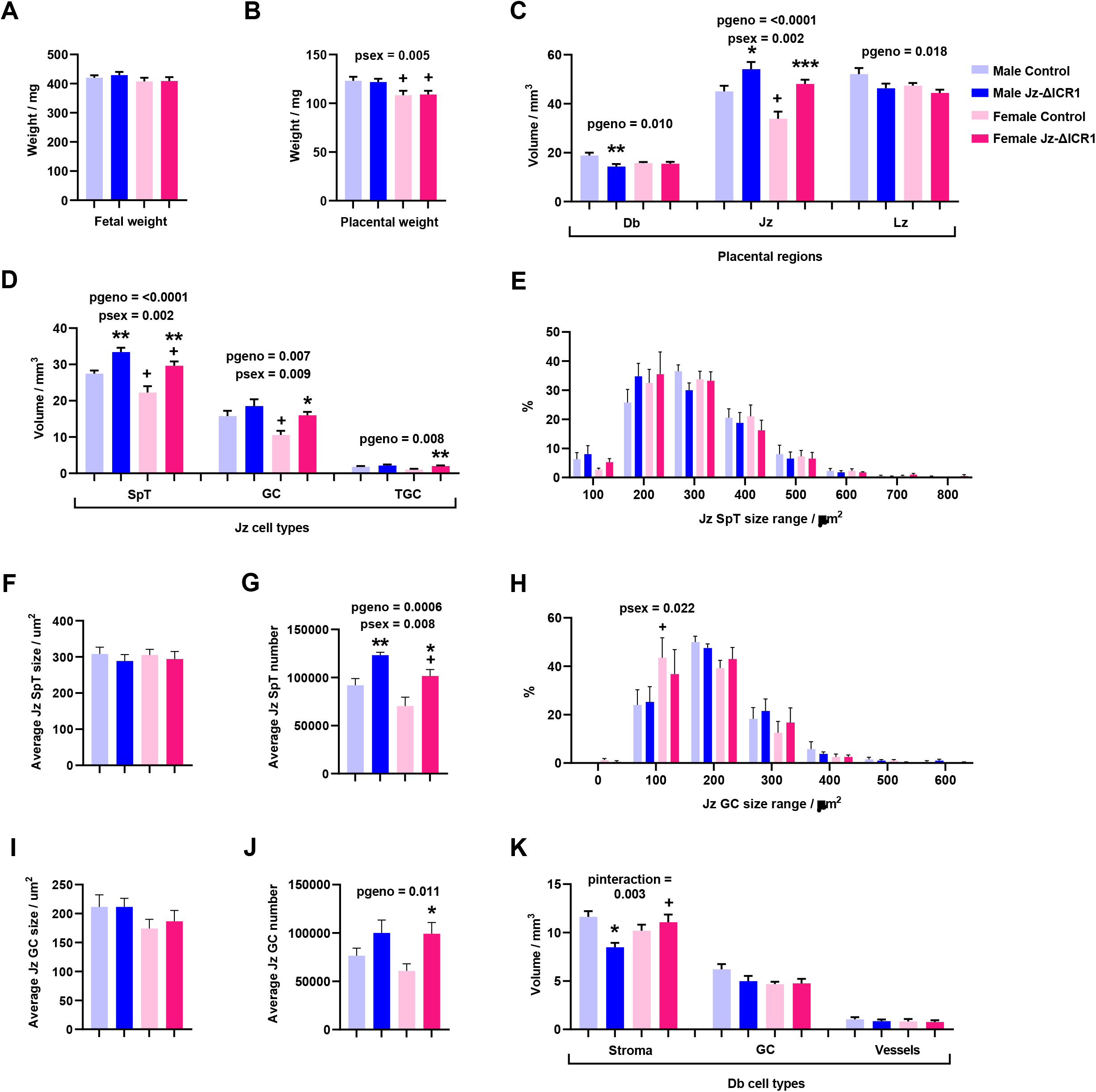
Jz-ΔICR1 increases the formation of endocrine cells in the Jz of the mouse placenta. (A) Fetal weight and (B) placental weight of D16 males (Control n = 29 and Jz-ΔICR1 n = 27) and females (Control n = 17 and Jz-ΔICR1 n = 27) from individual pups, across 13 litters. Volume of (C) placental regions and (D) Jz cell types (n = 8 per genotype/sex, across 7 litters). Jz SpT cell (E) size distribution, (F) average size and (G) average number, and Jz GC cell (H) size distribution, (I) average size and (J) average number (n = 4 per genotype/sex, across 4 litters). (K) Volume of Db cell types (n = 8 per genotype/sex, across 7 litters). Values presented as mean + SEM with significance assessed by two-way ANOVA and pairwise T-test (pgenotype < 0.05 = * < 0.01 = **, < 0.001 = ***, psex < 0.05 = +). Decidua (Db), Junctional zone (Jz), Labyrinth zone (Lz), spongiotrophoblast (SpT), glycogen cell (GC), trophoblast giant cell (TGC).

Further stereological analysis of the placenta revealed that increased Jz volume with Jz-ΔICR1 was attributable to increased volume of SpT (+33%), GC (+51%) and TGCs (+96%) in females and increased volume of SpT (+22%) in males (Fig. 3D). The total volume of SpT in both genotypes and total volume of GC in controls was less in female placentas compared to males. The distribution and average cell size of SpT cells in the Jz was unaffected by genotype (Fig. 3E and F), however, the average number of SpT cells was increased by 44% in females and 34% in males in response to Jz-ΔICR1 (Fig. 3G). The average number of SpT cells in Jz-ΔICR1 placentas was less in females compared to males (Fig. 3G). The distribution and average cell size of Jz GC cells was also unaffected by genotype (Fig. 3H and I), although there was a greater percentage of GC that were within the 100 μm^2^ size range in control females versus control males (Fig. 3H). There was also a 63% increase in the average number of Jz GC cells in female placentas with Jz-ΔICR1, but no significant effect observed in males (Fig. 3J). There was an interaction between genotype and sex in determining the total volume of decidual stroma (Db_S); whereby Db_S volume was reduced by Jz-ΔICR1 in male, but not female conceptuses, and Db_S volume was greater in Jz-ΔICR1 females compared to Jz-ΔICR1 males (Fig. 3K). There was no effect of genotype or sex on the volume of GC and vessels in the decidua.

As indicated by Ki67 staining, there was an overall >3-fold increase in cell proliferation in the Jz of Jz-ΔICR1 conceptuses, an effect significant in all three Jz cell types (Fig. 4B). In contrast, as determined by cleaved caspase-3 staining, there was no effect of Jz-ΔICR1 on the number of cells undergoing apoptosis in the Jz (Fig. 4D). There was no effect of fetal sex on Jz cell proliferation or apoptosis. Representative negative control images can be found in Fig S1C,D.

**Fig. 4.**
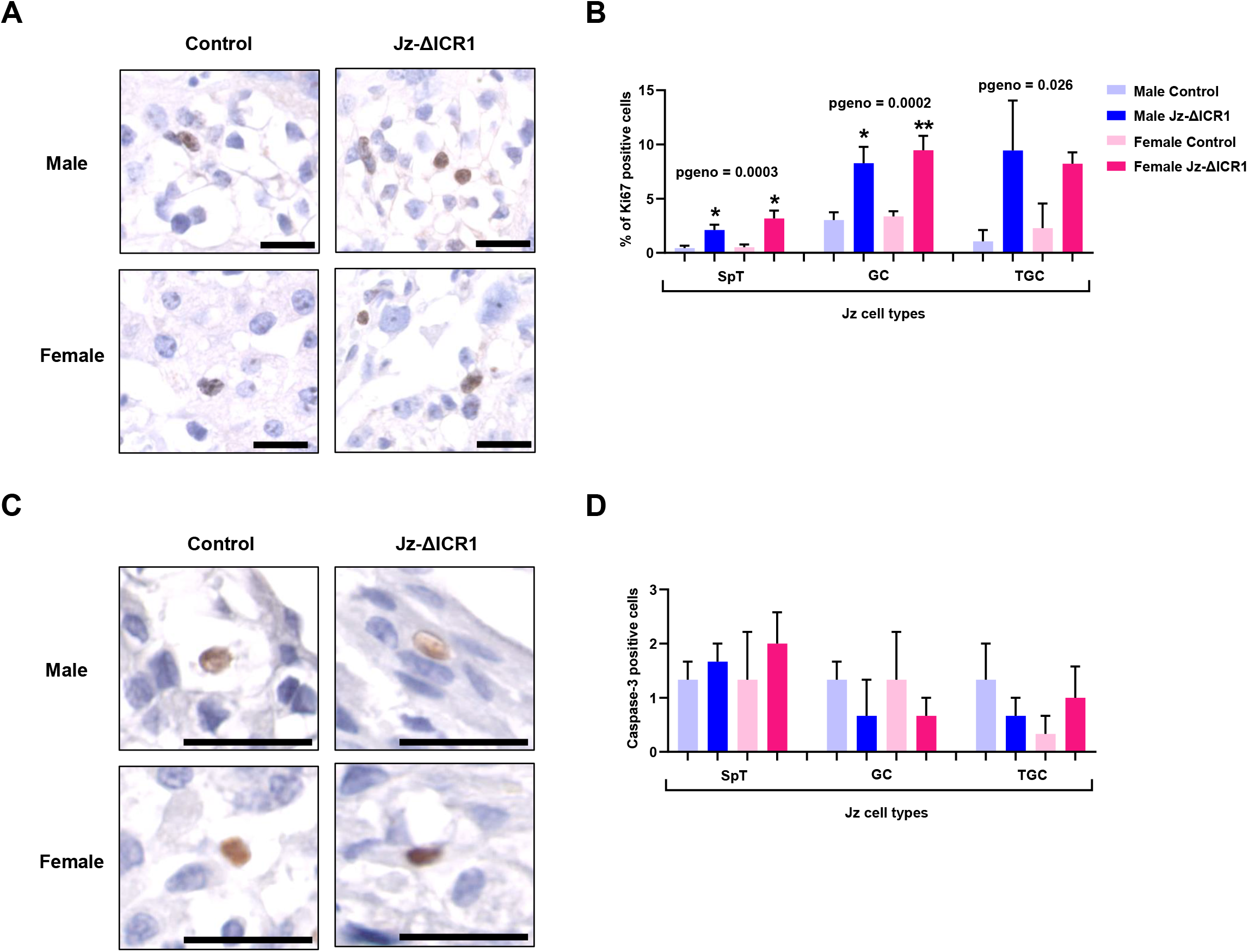
Jz-ΔICR1 increases proliferation Jz cell types but does not alter apoptosis levels in the Jz of the mouse placenta. Immunostaining for (A) Ki67 in male and female placentas with (B) the percentage of Ki67 positive cells in the Jz, and immunostaining for (C) cleaved caspase-3 in male and female placentas with (D) the number of cleaved caspase-3 positive cells in the Jz. Data were obtained on D16. The black scale bar represents 25 μm. Values presented as mean + SEM with n = 3-5 per genotype/sex, across 5 litters. Significance assessed by two-way ANOVA and pairwise T-test (pgenotype < 0.05 = *, < 0.01 = **). Spongiotrophoblast (SpT), glycogen cell (GC), trophoblast giant cell (TGC).

### Jz-ΔICR1 did not affect the expression of Jz cell markers but increased total placental glycogen storage in females

Whilst total volumes of the three Jz lineages were increased, expression of the SpT marker *Prl8a8,* the GC markers *Gjb3* and *Pcdh12,* and the TGC marker *Hand1* were all unaffected by Jz-ΔICR1 at the cellular level regardless of fetal sex (Fig. 5A). However, expression of *Gjb3, Pcdh12* and *Hand1* was lower in the placental Jz from females compared to males, and in the case of *Gjb3* and *Pcdh12,* pairwise comparisons revealed this was significant for controls only (Fig. 5A). We next investigated whether the increased GC volume in Jz-ΔICR1 placentas impacted placental glycogen metabolism. Jz-ΔICR1 did not affect the expression of the glucose transporter *Slc2a1* or key glycogen synthesis pathway genes in the placental Jz (Fig. 5B). However, overall, *Gys1* and *Gbe1* were more highly expressed in the placentae of males compared to females (Fig. 5B). Placental glycogen concentration was also not altered by Jz-ΔICR1 (Fig. 5C). However, total placental glycogen was increased by Jz-ΔICR1, an effect that was significant in females (+34%) when data were separated by sex (Fig. 5D). Visualisation of GC by PAS staining did not identify any overt differences in the spatial localisation of GC in the placenta in response to Jz-ΔICR1 (Fig. 5E). We further assessed the integrity of the Jz/Lz and Db/Jz boundaries by *in situ* hybridisation for the Jz marker *Tpbpa* and the SpT marker *Prl8a8* and qualitative assessment revealed no overt differences between control and Jz-ΔICR1 placentas (Fig. 5F,G).

**Fig. 5.**
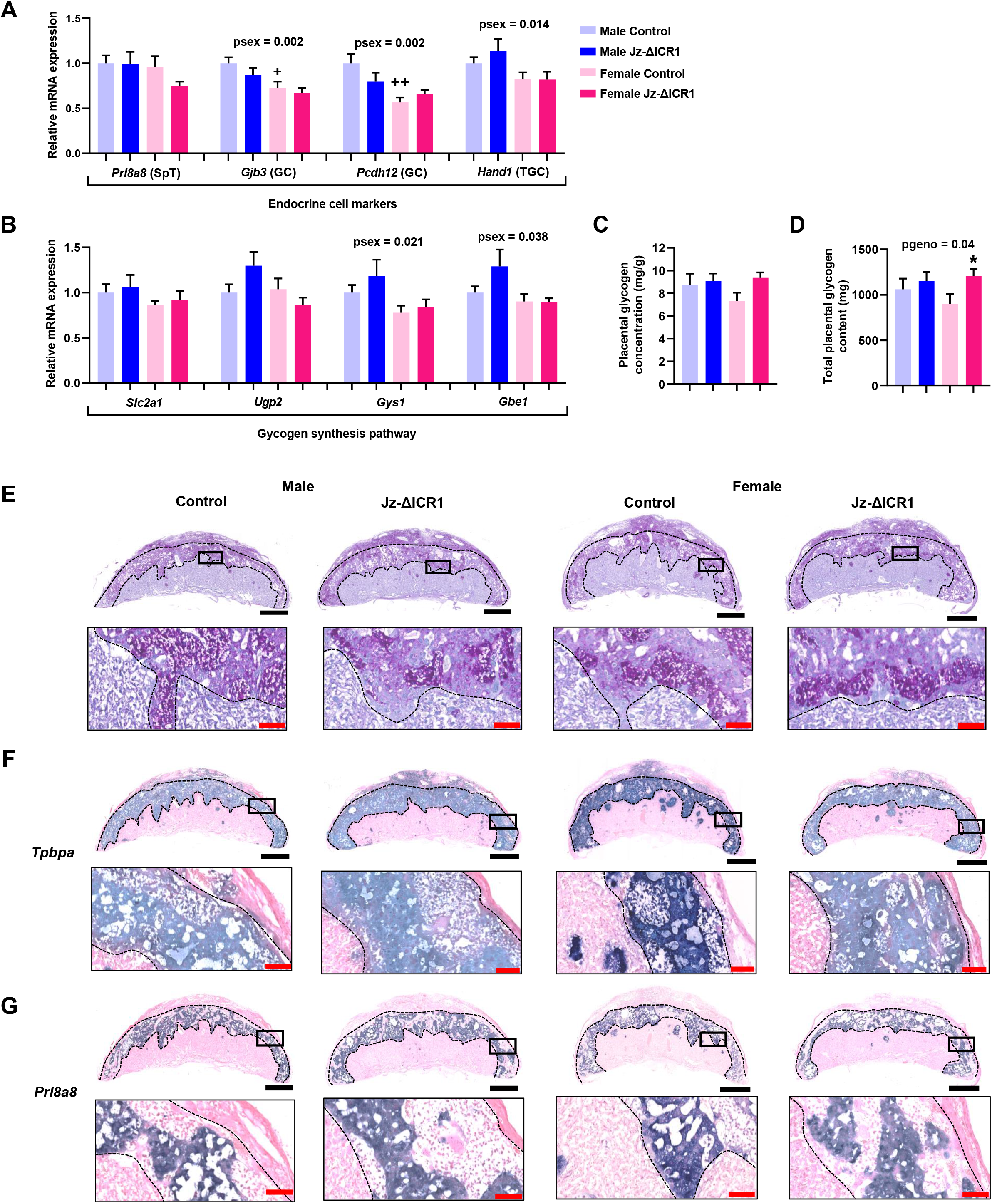
Jz-ΔICR1 increases total placental glycogen content in females, but does not alter placental glycogen concentration or the expression of glycogen synthesis pathway genes in mice. Jz expression of (A) endocrine cell markers and (B) glycogen synthesis pathway genes in Jz samples relative to the geometric mean of housekeeping genes (*Hprt* and *Ywhaz*) using qPCR (n = 8-10 per genotype/sex, across 11 litters). (C) Placental glycogen concentration (mg/g) and (D) total placental glycogen content (mg) in males and females with Jz-ΔICR1 (n = 8 per genotype/sex, across 8 litters). Data were obtained on D16. Values presented as mean + SEM with significance assessed by two-way ANOVA and pairwise T-test (psex < 0.05 = +, < 0.01 = ++). (E) PAS stain of glycogen containing cells and *in situ* hybridization of (F) *Tpbpa* and (G) *Prl8a8* in males and females. Black boxes represent the area magnified in the image below. Black bar represents 1 mm. Red bar represents 100 μm.

### Jz-ΔICR1 results in increased expression of the Jz hormone *Psg23*

We next investigated whether placental endocrine function was affected by Jz-ΔICR1. The expression of the steroidogenic pathway genes *Hmgcr, Stard1, Cyp11a1, Hsd3b1* and *Cyp17a1* were unaltered by Jz-ΔICR1, although *Stard1* was expressed at a lower level in the placental Jz of females compared with males (Fig. 6A). Similarly, expression of members of the *Prl* gene family *Prl2c2, Prl3b1, Prl3d1, Prl6a1* and *Prl7b1* and the angiogenic regulators *Flt1* and *Vegfa* were unaffected by Jz-ΔICR1, although expression of *Prl2c2* and *Prl3d1* by the placental Jz was lower in females compared to males (Fig. 6B and C). Whilst expression of *Psg21* was unaffected by Jz-ΔICR1, we observed a >2-fold increase in expression of *Psg23* in the Jz from both males (+2.3-fold) and females (+2.2-fold) (Fig. 6C). Overall, expression of *Psg23* was lower in placental Jz of females compared to males (Fig. 6C). *In situ* hybridisation revealed that in control placentas, *Psg*23 was localised to the Jz, with high levels of expression in the SpT and weak expression within GC. The spatial localisation of *Psg23* was maintained in Jz-ΔICR1 placentas, although staining intensity was notably greater when compared to controls (Fig. 6D).

**Fig. 6.**
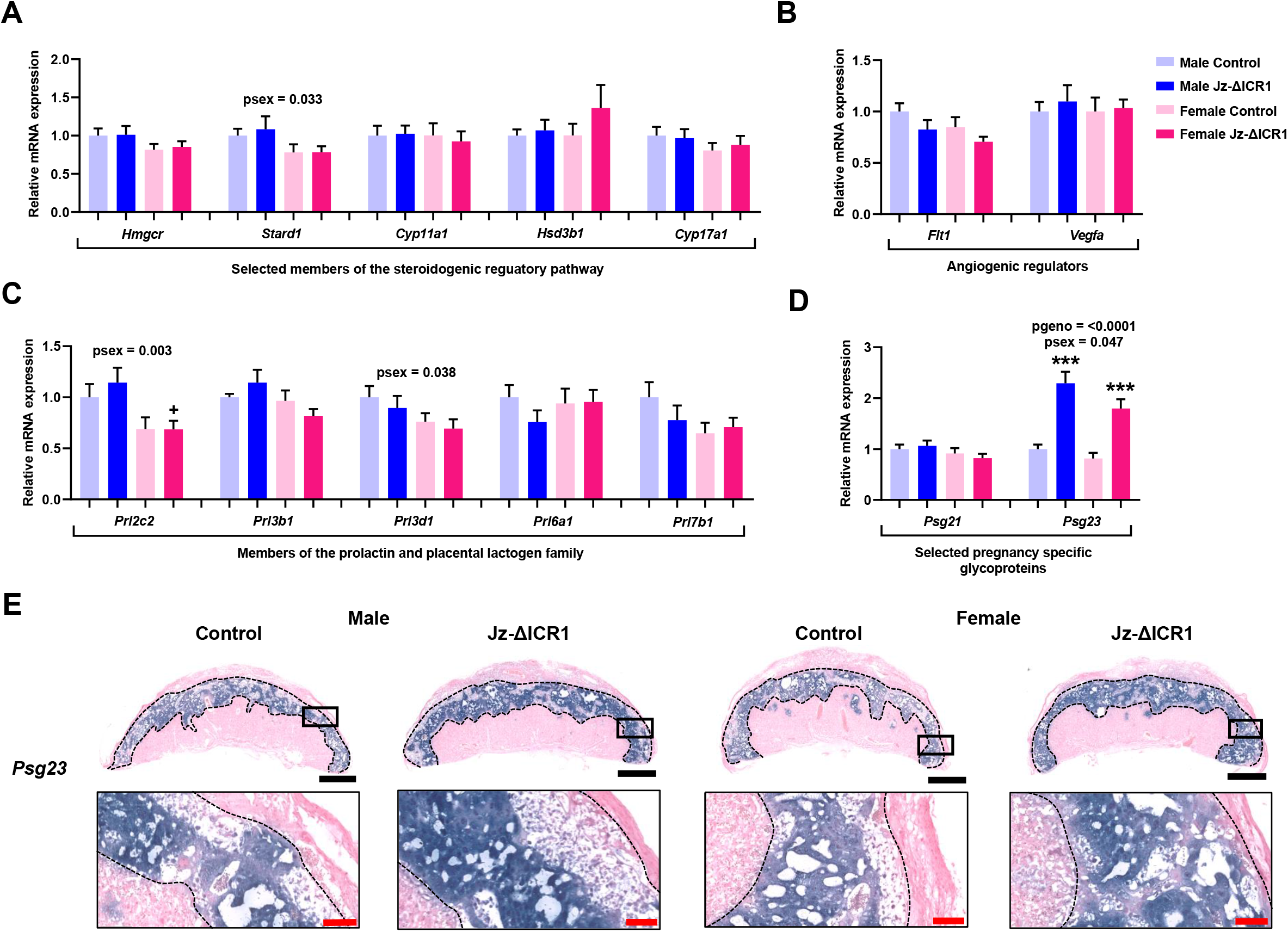
Jz-ΔICR1 alters the expression of *Psg23,* a Jz hormone, but not steroidogenic pathway or angiogenic regulatory genes. Expression of (A) steroidogenic pathway regulatory genes, (B) angiogenic regulatory genes, and selected mouse (C) *Prl* and (D) *Psg* genes in Jz samples relative to the geometric mean of housekeeping genes (*Hprt* and *Ywhaz*) using qPCR (n = 9-10 per genotype/sex, across 11 litters). Data were obtained on D16. Values presented as mean + SEM with significance assessed by two-way ANOVA and pairwise T-test (pgenotype < 0.001 = ***). (E) *In situ* hybridization of *Psg23* in D16 mouse placentas in males and females. Black boxes represent the area magnified in the image below. Black bar represents 1 mm. Red bar represents 100 μm.

### Jz-ΔICR1 alters the protein expression of IGF2 signalling factors

To inform on the mechanism through which loss of imprinting of the ICR1 domain (via Jz deletion of ICR1) regulates Jz development, we quantified the abundance of IGF2 receptors (IGF1R, IGF2R and INSR) and downstream members of the PI3K-AKT and MAPK pathways (PI3K subunits P85, P110α, P110ß, as well as phosphorylated (p) and total (T-) AKT, GSK3, P38 and MAPK 42/44) using western blotting (Fig. 7). In males, p/T-AKT (phosphorylated to total AKT), T-GSK3, pP38, p/T-P38 (phosphorylated to total P38) and pMAPK were significantly increased by Jz-ΔICR1, with a trend for an increase in P110β (p=0.07) (Fig. 7A,C). However, the levels of IGF1R, T-AKT and p/T-GSK3 (phosphorylated to total GSK3) were significantly decreased in the Jz of males with Jz-ΔICR1. Conversely, in females, there was a significant increase in INSR, P85, pAKT, T-AKT, with a trend for an increase in IGF2R (p=0.08) in the Jz in response to Jz-ΔICR1 (Fig. 7B,D). Females with Jz-ΔICR1 also had a significant decrease in the level of T-GSK3 and a trend for a decrease in T-MAPK (p=0.06), compared to controls. Changes in protein abundance with Jz-ΔICR1 were not associated with corresponding changes in gene expression as assessed by qPCR in either the placental Jz of male or female fetuses (Fig. S2). Although there was a significant effect of fetal sex on Jz expression of *Insr, Gsk3, N-ras* and *Mek1* (Fig. S2).

**Fig. 7.**
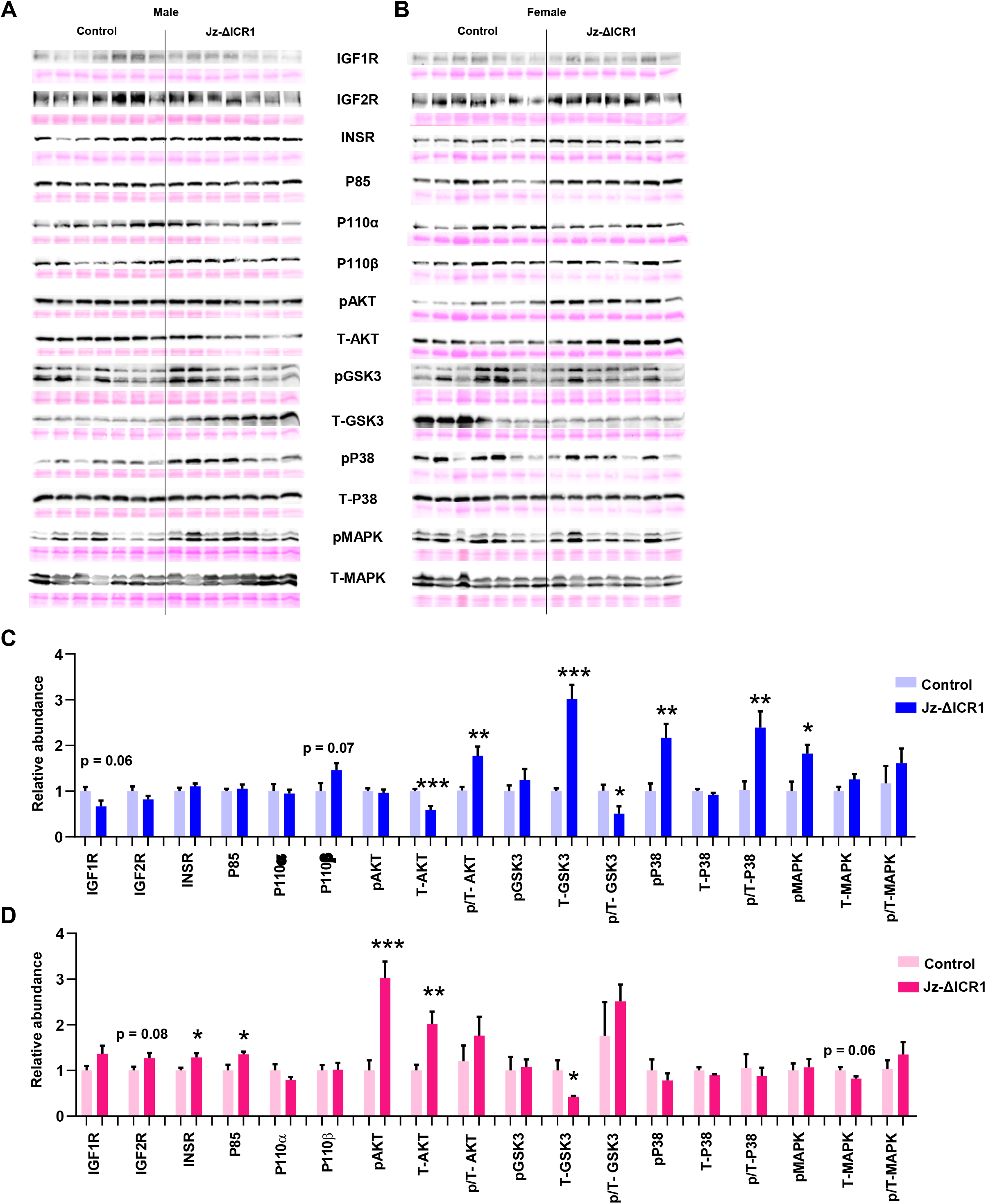
Jz-ΔICR1 alters the protein expression of IGF2 signalling factors and downstream members of the PI3K-AKT and MAPK pathway in the Jz. (A & B) Representative ponceau shown to indicate protein loading with the abundance, and (C & D) quantification of IGF2 signalling proteins in (A & C) males and (B & D) females. Data were obtained on D16 from n = 7 per sex genotype/sex, across 8 litters. Values presented as mean + SEM with significance assessed by T-test (p < 0.05 = *, p < 0.01 = **, p < 0.01 = ***)

### Other contributing mechanisms

Cre-mediated ablation of the transcription factor *Tfap2c* leads to a reduction in Jz size, with increased expression of *H19* and decreased expression of *Psg23* in isolated Jz (Sharma et al., 2016). Thus, we examined whether expression of *Tfap2c* was altered in response to Jz-ΔICR1 and could contribute to the placental Jz phenotype observed. However, there was no significant effect of Jz-ΔICR1 on Jz *Tfap2c* expression on D16 (Fig. S3). Overall, expression of *Tfap2c* by the placental Jz was ~20% lower in females compared to males.

In addition to maternal re-activation of *Igf2* and downregulation of *H19,* Jz-ΔICR1 will also result in expression of the normally-silenced maternal *mir-483* (located within intron 2 of *Igf2),* and reduced expression of *mir-675* (located within exon 1 of *H19).* Although no miR-483 targets have been identified, some for miR-675 have been reported, including *Igf1r, Rapgap1, Egr3* and *Slc44a1* (Keniry et al., 2012). However, the expression of these by the placental Jz was not significantly affected by Jz-ΔICR1 (Fig. S2A and Fig. S4). Although *Rapgap1* was expressed at a lower level in females compared to males, an effect significant by pairwise comparison for control fetuses (Fig. S4).

## Discussion

Emerging studies demonstrate a role for imprinted genes in regulating placental endocrine capacity (John, 2013; 2017), with constitutive gene manipulations showing that the maternally expressed genes *Ascl2* and *Phlda2* restrict Jz size (Tunster et al., 2015, 2016), and paternally expressed genes *Peg3* and *Igf2* appearing to enhance Jz size (Esquiliano et al., 2009; Tunster et al., 2018). Furthermore, we recently reported that conditional loss of *Igf2* in cells of the Jz affects placental endocrine capacity in a sexually dimorphic manner (Aykroyd et al., 2020). In the present study, we generated a novel loss of imprinting (LOI) model to investigate the role of imprinting of the ICR1 domain in modulating placental endocrine capacity. We utilised *Tpbpa*-Cre-mediated deletion of ICR1 to drive LOI of the ICR1 domain specifically in cells of the Jz of the mouse placenta (Jz-ΔICR1). Quantification of gene expression in isolated Jz samples revealed a ~30% increase in *Igf2* expression and a concomitant ~40% reduction in *H19* expression within the Jz following maternal transmission of the floxed allele. Although the magnitude of changes in *Igf2* and *H19* expression may partly reflect the presence of *Tpbpa*-negative cells in isolated Jz samples, a similar level of *Igf2* reactivation and *H19* suppression has been seen in another study involving a 1.6 kb deletion spanning ICR1 (Thorvaldsen et al., 1998). Moreover, our findings reinforce the idea that there may be additional mechanisms controlling the placental expression of these imprinted genes from the maternal allele (Kaffer et al., 2001; Nordin et al., 2014; Sasaki et al., 2000). Importantly, correct spatial localisation of *Igf2* and *H19* transcripts was maintained in Jz-ΔICR1 placentas, and there was no effect on expression of *Igf2* or *H19* in the Lz.

Whilst placental weight was unaffected, Jz-ΔICR1 resulted in a slight reduction in Lz and Db volumes with a concomitant increase in Jz volume in placentas of both male and female fetuses. When analysed independent of sex, the increased Jz volume was attributable to increased volumes of all three Jz constituent cell types (SpT, GC and TGC), with no effect on average size of SpT or GC. This is consistent with findings in the *H19*^Δ13^ model, in which cell size was unaltered (Esquiliano et al., 2009), and suggests that the increased volume of Jz cell types is related to an expansion in cell number. Whilst IGF2 may have an antiapoptotic function (Sferruzzi-Perri et al., 2017), the level of the apoptotic marker caspase-3 was unaltered in the Jz by Jz-ΔICR1. However, there was more than a 3-fold increase in the percentage of Jz cells positive for the proliferation marker Ki67 in Jz-ΔICR1 placentas compared with controls. In addition to the well-established role of IGF2 in promoting trophoblast proliferation (Chen et al., 2016; Forbes et al., 2008), *H19* has also been linked to regulating placental cell proliferation. A Jz specific manipulation that results in upregulated *H19* expression also led to a 75% reduction in the number of Ki67 positive cells compared to control placentas at day 12.5 (Sharma et al., 2016). Taken together, these data show that the enhancement of Jz growth observed with Jz-ΔICR1 is due to an increase in proliferation and not a change in Jz cell size or apoptosis, which occurs as result of both *Igf2* and *H19* mis-expression.

In addition to mediating the transfer of nutrients from mother to fetus, the placenta also stores glucose as glycogen (Lefebvre, 2012; Tunster et al., 2020). These placental glycogen stores are thought to provide an important source of energy to support fetal growth during late gestation (Coan et al., 2006). Whilst Jz-ΔICR1 did not affect expression of key glycogenesis pathway genes measured, or weight adjusted glycogen concentration within the Jz, total placental glycogen content was increased. These findings are consistent with prior work showing that manipulation of the *Igf2-H19* locus affects placental glycogen levels (Esquiliano et al., 2009; Lopez et al., 1996). The effect of Jz-ΔICR1 to increase total placental glycogen was most pronounced for female fetuses, and overall, reflected greater GC number. These data indicate that imprinting of ICR1 in the Jz acts, at least in part, to restrict placental glycogen storage, thus reducing the necessity of the mother to invest resources that she may otherwise require for maintenance of health.

Expression of markers of Jz cell lineages and various endocrine related genes were largely unaltered at the cellular level by Jz-ΔICR1. However, since the abundance of the cell types expressing these genes is increased, total placental expression/output of such endocrine mediators would be predicted to be increased. A notable exception to this was *Psg23* expression, which increased >2-fold in the placental Jz of both males and females in response to Jz-ΔICR1. PSGs are thought to have a predominantly immune-modulatory function, and PSG23 is one of the most abundant PSGs expressed in the mouse placenta in late gestation (McLellan et al., 2005). Human PSGs induce the production of antiinflammatory cytokines *in vitro* (Snyder et al., 2001). Additionally, murine PSG23 and human PSG1 share a common function in promoting feto-placental blood supply via induction of vascular remodelling and angiogenesis (Lisboa et al., 2011; Wu et al., 2008). Given the upregulation of *Psg23* expression, future work should evaluate whether maternal spiral artery remodelling, inflammatory cytokine production, placental vascularisation and utero-placental blood flow may be altered by Jz-ΔICR1. However, the expression of the angiogenesis regulators *Flt1* and *Vegfa* were unaffected by Jz-ΔICR1 in this study. Taken together, Jz-ΔICR1 appears to increase placental endocrine capacity and function through two mechanisms; firstly, by increasing the abundance of the endocrine cell lineages (SpT, GC and TGC), and secondly by driving increased expression of *Psg23* at the cellular level.

The mechanism underlying enhanced *Psg23* with Jz-ΔICR1 is unclear. Interestingly, *Tpbpa-* Cre-mediated ablation of the transcription factor *Tfap2c* resulted in a reduction in Jz size, with increased expression of *H19* and decreased expression of *Psg23* in the Jz (Sharma et al., 2016). However, expression of *Tfap2c* was unaltered in response to Jz-ΔICR1, which argues against a role for altered *Tfap2c* in our model. Maternal re-activation of *Igf2* and downregulation of *H19,* will also result in expression of the normally-silenced maternal *mir-483* (located within intron 2 of *Igf2),* and reduced expression of *mir-675* (located within exon 1 of *H19).* miRNAs can regulate gene expression and have been implicated in Jz cell proliferation and differentiation (Sharma et al., 2019). As miR-675 has been linked with placental growth suppression (Keniry et al., 2012), dysregulation of miR-483 or miR-675 could contribute to the phenotypes observed in response to Jz-ΔICR1. However, we did not observe any alterations in expression of several miR-675 targets, which include *Igfr1,* in response to Jz-ΔICR1. Furthermore, expression of *Psg23* is unaltered in the *H19^Δ3^* model (Keniry et al., 2012), suggesting that *Psg23* is not a target of miR-675. No miR-483 targets have been identified to date, thus the contribution of perturbed miR-483 expression to the phenotype observed with Jz-ΔICR1 requires study.

We wanted to further explore the underlying molecular mechanisms through which imprinting of ICR1 may exert a regulatory influence on placental endocrine capacity. Since IGF2 is elevated ~30% in the Jz as a result of Jz-ΔICR1, and the phenotype in the *H19*^Δ13^ model was attributed to elevated IGF2 (Leighton et al., 1995a), we focused our attention on the abundance of receptors that bind to, and signalling pathways downstream of, IGF2. In females, protein levels of INSR, PI3K-P85, phosphorylated AKT and total AKT were increased alongside decreased levels of total GSK3 in the placental Jz response to Jz-ΔICR1. As the PI3K-AKT signalling pathway inhibits GSK3, which is a negative regulator of glycogen synthesis (Cross et al., 1995; Diehl et al., 1998), our findings are consistent with increased placental glycogen synthesis in Jz-ΔICR1 females. Deletion of a negative regulator of PI3K-AKT signalling has been shown to increase placental Jz size in mice (Church et al., 2012). This signalling pathway is also implicated in the proliferation and differentiation of individual trophoblast cells in the placenta, notably GC and TGC (Kent et al., 2010; Lee at al., 2019; Sferruzzi-Perri et al., 2016). Thus, enhanced PI3K-AKT activation (via INSR) could also explain the increase in Jz formation in females with Jz-ΔICR1.

In males with Jz-ΔICR1, there was no increase in the abundance of signalling receptors for IGF2 (there was even a trend for reduced IGF1R), and abundance of total AKT was decreased and levels of total GSK3 increased in the placental Jz. However, there was a trend for elevated PI3K-P110β, and the ratio of active phosphorylated to total AKT was increased, whilst the ratio of inactive phosphorylated to total GSK3 increased in Jz-ΔICR1 males. These changes all suggest enhanced PI3K-AKT signalling. However, unlike in females, this enhanced PI3K-AKT signalling was not associated with an increase in total placental glycogen or GC and TGC abundance in Jz-ΔICR1 males. These data collectively suggest that the precise mechanism through which activation of PI3K-AKT signalling occurs, or the existence of other signalling pathways, are important for modulating glycogen levels and Jz morphogenesis in the female and male placenta. Indeed, males, but not females, showed an increased level of phosphorylated MAPK, phosphorylated P38 and phosphorylated to total P38 ratio in response to Jz-ΔICR1. Members of the MAPK pathway are activated by IGF2 to promote cell proliferation (Forbes and Westwood, 2008). P38 signalling also regulates programmed cell death pathways (Cheng and Feldman, 1998; Li et al., 2003) and is important for murine placental Jz formation, particularly SpT differentiation (Mudgett et al., 2000). While there was no change in apoptosis with Jz-ΔICR1, elevated abundance and activation of MAPK pathway could also explain the enhanced in Jz formation in males in response to Jz-ΔICR1. Changes in Jz protein abundance with Jz-ΔICR1 were not linked to alterations in gene expression, highlighting a role for post-transcriptional regulation in mediating the changes seen for males and female fetuses, which require study in further work.

Previous studies manipulating the expression of genes within the ICR1 domain have demonstrated profound effects on fetal growth. For instance, maternal inheritance of the *H19^Δ13^* allele, which deletes both *H19* and ICR1 and results in maternal reactivation of *Igf2,* causes a 27% increase in fetal weight (Leighton et al., 1995a). Similarly, maternal inheritance of a deletion spanning only the ICR1 results in a 17% increase in neonatal weight (Thorvaldsen et al., 1998), whereas targeted deletion of *H19* (*H19^Δ3^*) which results in partial (~25%) reactivation of maternal *Igf2,* enhanced fetal weight by ~8% (Ripoche et al., 1997). Despite a ~30% increase in *Igf2* expression in the Jz and the enhanced placenta endocrine and glycogen storage capacity, fetal weight was not increased in our Jz-ΔICR1 model. There are several potential explanations for this observation. Firstly, deletion of ICR1 was specifically targeted to cells of the Jz, and *Igf2* expression in the Lz or fetus (which are unaltered) may be relatively more important for fetal growth. This notion is in line with previous studies using manipulations that only affect one compartment of the placenta or fetus (Aykroyd et al., 2020; Sandovici et al., 2019; Sferruzzi-Perri et al., 2011). Secondly, we assessed impacts on fetal weight on day 16 of pregnancy only. This day was chosen as it is when the placental Jz is at its largest size (Coan et al., 2006) and mouse dams are most insulin resistant to favour fetal nutrient supply (Musial et al., 2016). Hence, work is required to explore if changes in placental endocrine phenotype with Jz-ΔICR1 offer benefit when the fetus enters its exponential growth phase in the lead up to term. Thirdly, we generated litters of mixed genotypes, such that both control and Jz-ICR1Δ conceptuses were exposed to the same *in utero* environment. Whilst this approach serves to normalise variations in the maternal environment, control littermates will be exposed to the potentially altered *in utero* environment caused by enhanced endocrine capacity of Jz-ICR1Δ placentas and may not show the normal pattern of fetal growth. Indeed, the phenotype of genetically wild-type littermates has been shown to be influenced by mutant littermates with phenotypes affecting placental endocrine function (López-Tello et al. 2019; Tunster et al. 2016). Future work may address any possible dilution of an effect on fetal growth by undertaking comparisons between litters comprised entirely of Jz-ΔICR1 or wild-type conceptuses.

In light of accumulating evidence of sexual dimorphism in placental adaptations to genetic and/or environmental perturbations (Aykroyd et al., 2020; Barke et al., 2019; Kalisch-Smith et al., 2017; Napso et al., 2019; Rosenfeld, 2015) and offspring outcomes (Christoforou and Sferruzzi-Perri, 2020; Dearden et al., 2018; Rodgers and Sferruzzi-Perri, 2021), we accounted for fetal sex in our analyses. Whilst Jz size was increased in placentas of both male and female conceptuses in response to Jz-ΔICR1, the underlying mechanisms appear to exhibit a degree of sexual dimorphism. For instance, the increase in Jz volume of Jz-ΔICR1 females was attributed to increased volumes of all three Jz cell types, whereas statistical significance was achieved only for SpT in Jz-ΔICR1 males. Furthermore, abundance of individual components of the PI3K-AKT and MAPK signalling pathways in the Jz with Jz-ΔICR1 was dependent on fetal sex. We also found that placental weight and Jz size were greater in males compared to females, alongside increased expression of several Jz lineage markers (*Gjb3, Pcdh12, Hand1*), glycogen synthesis genes (*Gys1, Gbe1, Gsk3*), growth regulators (*Insr, N-ras, Mek1, Tfap2c, Rapgap1*) and endocrine related genes (*Stard1, Prl2c2, Prl3d1, Psg23*), irrespective of genotype. Our findings further emphasise the need to control or account for fetal sex in studies of fetal and placental developmental physiology.

In summary, Jz-ΔICR1 enhances endocrine cell formation and Jz hormone expression. Whilst this phenotype is observed in both sexes, the signalling pathway response mechanisms attributable to these alterations are sexually-dimorphic. Moreover, the expansion of the Jz occurs at the expense of Lz size which is although not associated with a significant alteration in placental or fetal weight. The influence of altered placental phenotype with Jz-ΔICR1 on fetal and maternal physiology is yet to be determined. Studies of the human placenta have reported perturbations in the regulation of the *Igf2-H19* locus and alterations in the expression/abundance of IGF2 signalling factors and placental hormones in pregnancy complications including gestational diabetes, fetal growth restriction and large for gestational age (Laviola et al., 2005; Le et al., 2013; Liao et al., 2017; Ngala et al., 2017; Su et al., 2016). Therefore, this study may provide valuable insight for understanding the pathogenesis of human pregnancy conditions. In addition, the magnitude of changes in *Igf2* and *H19* expression seen in this study supports the notion that there may be other regulatory mechanisms controlling the expression of these genes from the maternal allele in the placenta. Finally, our study highlights that maternal and paternal imprinted genes may govern the allocation of resources to the fetus additionally via modulation of placental endocrine function in pregnancy.

## Materials and methods

### Maintenance of transgenic mice

The experiments for this study were approved by the University of Cambridge Animal Welfare and Ethical Review Body performed under the UK Home Office Animals (Scientific Procedures) Act 1986. Homozygous *TpbpaCre* males (Simmons et al., 2007) were mated to heterozygous ICR floxed females (LoxP sites surrounding the ICR; termed ICR1 Flox; Srivastava et al., 2000) to generate litters containing fetuses with control and Jz-ΔICR1 placentae (Fig. 1). The transgenic mice were maintained on a C57BL/6NCrl (Charles River, UK) background for >10 generations. Mice were housed under a 12:12h light/dark photocycle, 22°C air temperature and 21% oxygen saturation with access to water *ad libitum* and standard laboratory chow (Rat and Mouse No.3; Special Dietary Services, UK).

### Tissue collection

Dams were killed by cervical dislocation on day 16 of pregnancy (presence of a copulatory plug denoted day 1 and term occurs ~ day 20). Placentae and fetuses were dissected from the dam uterus and weighed. Placentae were bisected on the short axis, one half was separated into individual Jz and Lz, as described previously by Sferruzzi-Perri et al., 2009, snap-frozen in liquid nitrogen and stored at −80°C for either gene expression or western blotting analysis. The remaining placental half was kept whole and either snap frozen in liquid nitrogen and stored at −80°C for placental glycogen content analysis, or fixed in 4% paraformaldehyde, dehydrated, embedded into paraffin wax and sectioned exhaustively from the mid-line at 8 μm for histological analysis. Fetal tails were collected for PCR to establish the Flox genotype (FPrimer: 5’-CAGGCCTGTCCTCACCTGAAC-3’, RPrimer: 5’-GCCAGCTTGCCTTGGCAACCCCTT-3’) and sex with *Sry* genotyping (FPrimer: 5’-GTGGGTTCCTGTCCCACTGC-3’, RPrimer: 5’-GGCCATGTCAAGCGCCCCAT-3’ with a PCR autosomal gene control FPrimer: 5’-TGGTTGGCATTTTATCCCTAGAAC-3’, RPrimer: 5’-GCAACATGGCAACTGGAAACA-3’). Placentae with weights closest to the mean value for each litter were selected from each genotype group for analysis.

### Stereological analysis

Every 20^th^ paraffin-embedded placental section was stained with haematoxylin and eosin (n = 8 per genotype/sex, across 8 litters). Images of each placental section were captured at 40x magnification using a NanoZoomer 2.0-RS (Hamamatsu, JP). The gross structure of each placental zone (Db, Jz and Lz) and the proportion of cells in the Jz and Db were analysed using the newCAST System (Visiopharm, DK) as described by Aykroyd et al., 2020. The average size of Jz GC was estimated by using the freehand annotation tool in the NDP.view2 viewing software (Hamamatsu, JP) and measuring the area of 100 individual Jz GC in a mid-line placental section (n = 4 per genotype/sex, across 4 litters). The average number of SpT and GC in the placental Jz was determined by dividing the SpT and GC volume by average cell size. A qualitative assessment of Jz interdigitation into the Lz, and Jz boundary integrity with the Db and Lz was also performed (n = 4 per genotype/sex, across 4 litters).

### *In situ* hybridisation

The expression of *Igf2, H19, Tpbpa, Prl8a8* and *Psg23* were localised in placental sections using *in situ* hybridisation. Previous studies have described the generation of probes for *Igf2* and *Prl8a8* (Aykroyd et al., 2020) and *Tpbpa* (Lescisin et al., 1988). A 659bp region of *Psg23* was amplified by PCR from wild-type placental cDNA (primer sequences: 5’-GCTGTGACCCTCTTGACTCT-3’, 5’-AAATGCCTCTGCCCTGCTAT-3’), cloned into the pDrive vector system (Qiagen, DE), with linearised vector used as template for probe transcription. For H19, the template for probe transcription was generated by PCR amplification of a 405 bp fragment from wild type placental cDNA using primers incorporating a T3 (forward primer: 5’-AATTAACCCTCACTAAAGGGTTGTCGTAGAAGCCGTCTGT-3’) or T7 (reverse primer: 5’-TAATACGACTCACTATAGGGGACAGGAGGGAGATGATGAAGT-3’) RNA Polymerase binding site, as described by Langford et al., 2018. The amplicon was purified using the Monarch PCR and DNA Cleanup Kit (NEB), and 1 ug used as template for transcription of digoxigenin labelled riboprobes using the DIG RNA Labelling mix (Sigma).

Probe hybridisation was performed overnight at 60°C, as described by Rakoczy et al., 2017. Staining was developed using BM-Purple alkaline phosphatase substrate (Sigma) and sections were counterstained using Nuclear Fast Red (Sigma-Aldrich, US). DIG-labelled sense riboprobes with identical sequences to the target mRNA were used as negative controls.

### Immunohistochemistry

Apoptosis and proliferation levels were measured by immunostaining in dewaxed and rehydrated midline placental sections with cleaved caspase-3 (Asp175) (Cell Signalling, US, 9661; 1:200) and Ki67 (Abcam, UK, ab264429; 1:500). Sections were incubated with goat-anti-rabbit secondary antibody (Abcam, UK, ab6720; 1:1000), streptavidin-horseradish peroxidase (Rockland, US, S000-03, 1:250) and stained with 3,3’-Diaminobenzidine (Abcam, UK). Haematoxylin was used as a counterstain before dehydrating and mounting the sections. Caspase and Ki67 positive cells were identified and counted in the placental Jz using NDP.view2 viewing software (Hamamatsu, JP), (n = 3-5 per genotype/sex, across 5 litters). Negative controls were prepared by the omission of primary antibodies.

### Glycogen assay

Amyloglucosidase was used to indirectly measure glycogen content in bisected placental halves (n = 8 per genotype/sex, across 8 litters), as previously described (Sferruzzi-Perri et al., 2013b).

### Placental gene expression

Total RNA was extracted and 5 μg reverse transcribed from paired isolated Jz and Lz placental tissues (n = 8-10 per genotype/sex, across 11 litters) using the RNeasy Plus Mini Kit (Qiagen, DE) and the High-Capacity cDNA Reverse Transcription Kit minus RT inhibitor (Applied Biosystems, US), according to manufacturer’s instructions. Primer sequences were sourced from publications and reported previously (Aykroyd et al., 2020) or designed using NCBI Primer Blast and produced by Sigma-Aldrich (Table S1). Only primers which produced a PCR product of the desired size, correct sequence and with amplification efficiencies of >85% were used. Samples were measured in duplicate on a 7500 fast real-time PCR machine (Applied Biosystems, US) with MESA Blue SYBR (Eurogentec, BE) under the following conditions: 3 minutes at 95°C then 40 cycles of: 30s at 95°C, 30s at 57°C, 90s at 72°C. The cycle threshold expression values for each gene were normalised to the geometric mean of housekeeping genes *Hprt* and *Ywhaz* for Jz samples and *Hprt* and *Polr2a* for Lz samples. All reference genes were unaltered by genotype or sex. Fold change was calculated according to the 2^-ΔΔCT^ method (Livak and Schmittgen, 2001) and represents an estimation of fold change at the cellular level, since expression is assessed in the Jz or Lz in isolation.

### Placental Jz protein expression

Protein was extracted from approximately 50mg of placental Jz tissue (n = 7 per genotype/sex, across 9 litters) using RIPA buffer (Thermo Scientific, US) containing cOmplete Mini EDTA-free protease inhibitor cocktail mix (Roche, CH). The protein concentration of Jz lysates was determined using the Bicinchoninic Acid protein assay (Thermo Scientific, US). Lysates were diluted to 2.5μg/μl in lysis buffer and 1xSDS, resolved using SDS-PAGE and transferred onto 0.2 μm nitrocellulose membranes (Bio-Rad Laboratories, US). Even protein loading and successful protein transfer was confirmed using Ponceau S stain (Sigma Aldrich) prior to probing with primary antibodies (Table S2). Antirabbit secondary antibody tagged to horseradish peroxidase (NA934 Cytiva, US, 1:10000 in 1xTBST with 2.5% milk/BSA) was used for all membranes. Bands were visualised using Scientific SuperSignal West Femto enhanced chemiluminescence (ECL) substrate (Thermo Scientific, US) and imaged using an iBright 1500 Imaging System (Invitrogen, US). Abundance of proteins were quantified using ImageJ analysis software (National Institutes of Health, US) to measure the pixel intensity of protein bands. Protein loading was controlled for by normalising against Ponceau S staining.

### Statistics

Prior to statistical analysis, a Prisms Grubbs’ test (GraphPad Software Inc.) was performed on all data sets to identify any outliers as a quality check. In the majority of cases, entire datasets did not contain outliers, and if they did, at most a single sample from a group was excluded. Final sample numbers are detailed within each table or figure legend. With the exception of protein abundance analyses, all data were analysed by two-way ANOVA (genotype and sex). If an overall significant effect of sex or genotype was identified, then planned comparisons using two-tailed T-tests were performed. Protein abundance was assessed for males and females separately to maintain a high sample size (n) per group and effect of genotype determined using two-tailed T-tests. Prism (GraphPad Software Inc., US) was used to perform statistical analyses with a significance value of p<0.05. Results are shown as mean +/± SEM, n = number of fetuses or placentae in each group.

## Acknowledgements

We are grateful to Miguel Constância and Ionel Sandovici for providing the *TpbpaCre* and ICR1 Flox animals used in this study and staff of the Combined Animal Facility for assistance in animal husbandry.

---

**Fig. S1. Negative controls for *in situ* hybridization and immunohistochemistry experiments in D16 mouse placentas**. Sense *in situ* hybridization probes for (A) *H19* and (B) *Psg23.* Secondary antibody only control used in immunohistochemistry experiments for (C) cleaved caspase-3 and (D) Ki67. Black boxes represent the area magnified in the image below. Black bar represents 1 mm. Red bar represents 100 μm.

**Fig. S2. Jz-ΔICR1 does not alter the gene expression of IGF2 signalling factors or downstream members of the PI3K-AKT and MAPK pathways.** Expression of (A) IGF2 signalling factors, (B) PI3K-AKT pathway genes and (C) RAS-MAPK-ERK pathway genes in Jz samples relative to the geometric mean of housekeeping genes (*Hprt* and *Ywhaz*) using qPCR (n = 8-10 per genotype/sex, across 11 litters). Data were obtained on D16. Values presented as mean + SEM with significance assessed by two-way ANOVA and pairwise T-test (psex < 0.05 = +).

**Fig. S3. Jz-ΔICR1 does not alter the gene expression of Tfap2c.** (A) Expression of Tfap2c in Jz samples relative to the geometric mean of housekeeping genes (Hprt and Ywhaz) using qPCR (n = 9-10 per genotype/sex, across 11 litters). Data were obtained on D16. Values presented as mean + SEM with significance assessed by two-way ANOVA and pairwise T-test (psex < 0.05 = +).

**Fig. S4. Jz-ΔICR1 does not alter the gene expression of miR-675 targets.** (A) Expression of miR-675 target genes in Jz samples relative to the geometric mean of housekeeping genes (*Hprt* and *Ywhaz*) using qPCR (n = 8-10 per genotype/sex, across 11 litters). Data were obtained on D16. Values presented as mean + SEM with significance assessed by two-way ANOVA and pairwise T-test (psex < 0.05 = +).

## References

Ahmed-Sorour, H. and Bailey, C. J. (1980). Role of ovarian hormones in the long-term control of glucose homeostasis. Interaction with insulin, glucagon and epinephrine. Horm. Res. 13, 396–403.

Angiolini, E., Fowden, A., Coan, P., Sandovici, I., Smith, P., Dean, W., Burton, G., Tycko, B., Reik, W., Sibley, C., et al. (2006). Regulation of placental efficiency for nutrient transport by imprinted genes. Placenta. 27, 98–102.

Angiolini, E., Coan, P.M., Sandovici, I., Iwajomo, O.H., Peck, G., Burton, G.J., Sibley, C. P., Reik, W., Fowden, A.L. and Constância, M. (2011). Developmental adaptations to increased fetal nutrient demand in mouse genetic models of Igf2-mediated overgrowth. The FASEB Journal. 25, 1737–1745.

Aykroyd, B.R., Tunster, S.J. and Sferruzzi-Perri, A.N. (2020). Igf2 deletion alters mouse placenta endocrine capacity in a sexually dimorphic manner. Journal of Endocrinology. 246, 93–108.

Baker, J., Liu, J.P., Robertson, E.J. and Efstratiadis, A. (1993). Role of insulin-like growth factors in embryonic and postnatal growth. Cell. 75, 73–82.

Barke, T.L., Money, K.M., Du, L., Serezani, A., Gannon, M., Mirnics, K. and Aronoff, D. M. (2019). Sex modifies placental gene expression in response to metabolic and inflammatory stress. Placenta. 78, 1–9.

Barlow, D.P., Stöger, R., Herrmann, B.G., Saito, K. and Schweifer, N. (1991). The mouse insulin-like growth factor type-2 receptor is imprinted and closely linked to the Tme locus. Nature. 349, 84–87.

Barton, S.C., Surani, M.A.H. and Norris, M.L. (1984). Role of paternal and maternal genomes in mouse development. Nature. 311, 374–376.

Bell, A.C. and Felsenfeld, G. (2000). Methylation of a CTCF-dependent boundary controls imprinted expression of the Igf2 gene. Nature. 405, 482–485.

Bell, A.C., West, A.G. and Felsenfeld, G. (1999). The protein CTCF is required for the enhancer blocking activity of vertebrate insulators. Cell. 98, 387–396.

Blois, S.M., Tirado-González, I., Wu, J., Barrientos, G., Johnson, B., Warren, J., Freitag, N., Klapp, B.F., Irmak, S., Ergun, S., et al. (2012). Early expression of pregnancy-specific glycoprotein 22 (PSG22) by trophoblast cells modulates angiogenesis in mice. Biology of reproduction. 86, 191.

Brelje, T.C., Stout, L.E., Bhagroo, N.V. and Sorenson, R.L. (2004). Distinctive roles for prolactin and growth hormone in the activation of signal transducer and activator of transcription 5 in pancreatic islets of langerhans. Endocrinology. 145, 4162–4175.

Burton, G.J. and Fowden, A.L. (2015). The placenta: a multifaceted, transient organ. Philosophical Transactions of the Royal Society B: Biological Sciences. 370, 20140066.

Camm, E.J., Botting, K.J. and Sferruzzi-Perri, A.N. (2018). Near to one’s heart: the intimate relationship between the placenta and fetal heart. Frontiers in physiology. 9, 629.

Cattanach, B.M. and Kirk, M. (1985). Differential activity of maternally and paternally derived chromosome regions in mice. Nature. 315, 496–498.

Cheng, H.L. and Feldman, E.L. (1998). Bidirectional regulation of p38 kinase and c-Jun N-terminal protein kinase by insulin-like growth factor-I. Journal of Biological Chemistry. 273, 14560–14565.

Christoforou, E.R. and Sferruzzi-Perri, A.N. (2020). Molecular mechanisms governing offspring metabolic programming in rodent models of in utero stress. Cellular and Molecular Life Sciences. 77, 4861–4898.

Church, D.N., Phillips, B.R., Stuckey, D.J., Barnes, D.J., Buffa, F.M., Manek, S., Clarke, K., Harris, A.L., Carter, E.J. and Hassan, A.B. (2012). Igf2 ligand dependency of Pten+/-developmental and tumour phenotypes in the mouse. Oncogene. 31, 3635–3646.

Coan, P.M., Burton, G.J. and Ferguson-Smith, A.C. (2005). Imprinted Genes in the Placenta--A Review. Placenta. 26, S10–S20.

Coan, P.M., Conroy, N., Burton, G.J. and Ferguson-Smith, A.C. (2006). Origin and characteristics of glycogen cells in the developing murine placenta. Developmental dynamics: an official publication of the American Association of Anatomists. 235, 3280–3294.

Coan, P.M., Fowden, A.L., Constancia, M., Ferguson-Smith, A.C., Burton, G.J. and Sibley, C.P. (2008). Disproportional effects of Igf2 knockout on placental morphology and diffusional exchange characteristics in the mouse. The Journal of physiology. 586, 5023–5032.

Constância, M., Hemberger, M., Hughes, J., Dean, W., Ferguson-Smith, A., Fundele, R., Stewart, F., Kelsey, G., Fowden, A., Sibley, C., et al. (2002). Placental-specific IGF-II is a major modulator of placental and fetal growth. Nature. 417, 945–948.

Constância, M., Angiolini, E., Sandovici, I., Smith, P., Smith, R., Kelsey, G., Dean, W., Ferguson-Smith, A., Sibley, C.P., Reik, W., et al. (2005). Adaptation of nutrient supply to fetal demand in the mouse involves interaction between the Igf2 gene and placental transporter systems. Proceedings of the National Academy of Sciences. 102, 19219–19224.

Cross, D.A., Alessi, D.R., Cohen, P., Andjelkovich, M. and Hemmings, B.A. (1995). Inhibition of glycogen synthase kinase-3 by insulin mediated by protein kinase B. Nature. 378, 785–789.

Czech, M.P. (1989). Signal transmission by the insulin-like growth factors. Cell. 59, 235–238.

Dearden, L. and Ockleford, C. (1993) Structure of human trophoblasts: correlation with function. In Biology of trophoblast (ed. Y. W. Loke and A. Whyte), pp 69–110. Elsevier, New York.

Dearden, L., Bouret, S.G. and Ozanne, S.E. (2018). Sex and gender differences in developmental programming of metabolism. Molecular metabolism. 15, 8–19.

DeChiara, T.M., Efstratiadis, A. and Robertsen, E.J. (1990). A growth-deficiency phenotype in heterozygous mice carrying an insulin-like growth factor II gene disrupted by targeting. Nature. 345, 78–80.

DeChiara, T.M., Robertson, E.J. and Efstratiadis, A. (1991). Parental imprinting of the mouse insulin-like growth factor II gene. Cell. 64, 849–859.

Diehl, J.A., Cheng, M., Roussel, M.F. and Sherr, C.J. (1998). Glycogen synthase kinase-3ß regulates cyclin D1 proteolysis and subcellular localization. Genes & development. 12, 3499–3511.

Edwards, C.A., Takahashi, N., Corish, J.A. and Ferguson-Smith, A.C. (2019). The origins of genomic imprinting in mammals. Reproduction, Fertility and Development. 31, 1203–1218.

Engel, N., Thorvaldsen, J.L. and Bartolomei, M.S. (2006). CTCF binding sites promote transcription initiation and prevent DNA methylation on the maternal allele at the imprinted H19/Igf2 locus. Human molecular genetics. 15, 2945–2954.

Esquiliano, D.R., Guo, W., Liang, L., Dikkes, P. and Lopez, M.F. (2009). Placental glycogen stores are increased in mice with H19 null mutations but not in those with insulin or IGF type 1 receptor mutations. Placenta. 30, 693–699.

Ferguson-Smith, A.C. (2011). Genomic imprinting: the emergence of an epigenetic paradigm. Nature Reviews Genetics. 12, 565–575.

Ferguson-Smith, A.C. and Surani, M.A. (2001). Imprinting and the epigenetic asymmetry between parental genomes. Science. 293, 1086–1089.

Ferguson-Smith, A.C., Sasaki, H., Cattanach, B.M. and Surani, M.A. (1993). Parental-origin-specific epigenetic modification of the mouse H19 gene. Nature. 362, 751–755.

Forbes, K. and Westwood, M. (2008). The IGF axis and placental function. Hormone Research in Paediatrics. 69, 129–137.

Fowden, A.L., Giussani, D.A. and Forhead, A.J. (2006). Intrauterine Programming of Physiological Systems: Causes and Consequences. Physiology. 21, 29–37.

Georgiades, P., Ferguson-Smith, A.C. and Burton, G.J. (2002). Comparative developmental anatomy of the murine and human definitive placentae. Placenta. 23, 3–19.

Gluckman, P.D., Hanson, M.A., Cooper, C. and Thornburg, K.L. (2008). Effect of in utero and early-life conditions on adult health and disease. New England Journal of Medicine. 359, 61–73.

Han, V.K., Hill, D.J., Strain, A.J., Towle, A.C., Lauder, J.M., Underwood, L.E. and D’Ercole, A. J. (1987). Identification of somatomedin/insulin-like growth factor immunoreactive cells in the human fetus. Pediatric Research. 22, 245–249.

Han, V.K., Lund, P.K., Lee, D.C. and D’Ercole, A.J. (1988). Expression of somatomedin/insulin-like Growth Factor Messenger Ribonucleic Acids in the Human Fetus: Identification, Characterization, and Tissue Distribution. The Journal of clinical endocrinology and metabolism. 66, 422–429.

Huang, C., Snider, F. and Cross, J.C. (2009). Prolactin receptor is required for normal glucose homeostasis and modulation of β-cell mass during pregnancy. Endocrinology. 150, 1618–1626.

Huang, X., Liu, G., Guo, J. and Su, Z. (2018). The PI3K/AKT pathway in obesity and type 2 diabetes. International journal of biological sciences. 14, 1483–1496.

Jarrett II, J.C., Ballejo, G., Saleem, T.H., Tsibris, J.C. and Spellacy, W.N. (1984). The effect of prolactin and relaxin on insulin binding by adipocytes from pregnant women. American journal of obstetrics and gynecology. 149, 250–255.

John, R.M. (2013). Epigenetic regulation of placental endocrine lineages and complications of pregnancy. Biochemical Society Transactions. 41, 701–709.

John, R.M. (2017). Imprinted genes and the regulation of placental endocrine function: Pregnancy and beyond. Placenta. 56, 86–90.

Jones, J.I. and Clemmons, D.R. (1995). Insulin-like growth factors and their binding proteins: biological actions. Endocrine reviews. 16, 3–34.

Kaffer, C.R., Grinberg, A. and Pfeifer, K. (2001). Regulatory mechanisms at the mouseIgf2/H19 locus. Molecular and cellular biology. 21, 8189–8196.

Kalisch-Smith, J.I., Simmons, D.G., Dickinson, H. and Moritz, K.M. (2017). Sexual dimorphism in the formation, function and adaptation of the placenta. Placenta. 54, 10–16.

Kanduri, C., Pant, V., Loukinov, D., Pugacheva, E., Qi, C.F., Wolffe, A., Ohlsson, R. and Lobanenkov, V.V. (2000). Functional association of CTCF with the insulator upstream of the H19 gene is parent of origin-specific and methylation-sensitive. Current Biology. 10, 853–856.

Kaneko-Ishino, T. and Ishino, F. (2019). Evolution of viviparity in mammals: what genomic imprinting tells us about mammalian placental evolution. Reproduction, Fertility and Development. 31, 1219–1227.

Keniry, A., Oxley, D., Monnier, P., Kyba, M., Dandolo, L., Smits, G. and Reik, W. (2012). The H19 lincRNA is a developmental reservoir of miR-675 that suppresses growth and Igf1r. Nature cell biology. 14, 659–665.

Kent, L.N., Konno, T. and Soares, M.J. (2010). Phosphatidylinositol 3 kinase modulation of trophoblast cell differentiation. BMC developmental biology. 10, 1–18.

Khatib, H. (2007). Is it genomic imprinting or preferential expression?. Bioessays. 29, 1022–1028.

Langford, M.B., Outhwaite, J.E., Hughes, M., Natale, D.R. and Simmons, D.G. (2018). Deletion of the Syncytin A receptor Ly6e impairs syncytiotrophoblast fusion and placental morphogenesis causing embryonic lethality in mice. Scientific reports. 8, 1–11.

Lau, M.M., Stewart, C.E., Liu, Z., Bhatt, H., Rotwein, P. and Stewart, C.L. (1994). Loss of the imprinted IGF2/cation-independent mannose 6-phosphate receptor results in fetal overgrowth and perinatal lethality. Genes & Development. 8, 2953–2963.

Laviola, L., Perrini, S., Belsanti, G., Natalicchio, A., Montrone, C., Leonardini, A., Vimercati, A., Scioscia, M., Selvaggi, L., Giorgino, R., et al. (2005). Intrauterine growth restriction in humans is associated with abnormalities in placental insulin-like growth factor signaling. Endocrinology. 146, 1498–1505.

LaVoie, H.A. and King, S.R. (2009). Transcriptional regulation of steroidogenic genes: STARD1, CYP11A1 and HSD3B. Experimental biology and medicine. 234, 880–907.

Le, T.N., Elsea, S.H., Romero, R., Chaiworapongsa, T. and Francis, G.L. (2013). Prolactin receptor gene polymorphisms are associated with gestational diabetes. Genetic testing and molecular biomarkers. 17, 567–571.

Lee, C.Q., Bailey, A., Lopez-Tello, J., Sferruzzi-Perri, A.N., Okkenhaug, K., Moffett, A., Rossant, J. and Hemberger, M. (2019). Inhibition of Phosphoinositide-3-Kinase Signaling Promotes the Stem Cell State of Trophoblast. Stem Cells. 37, 1307–1318.

Lefebvre, L. (2012). The placental imprintome and imprinted gene function in the trophoblast glycogen cell lineage. Reproductive biomedicine online. 25, 44–57.

Leighton, P.A., Ingram, R.S., Eggenschwiler, J., Efstratiadis, A. and Tilghman, S.M. (1995a). Disruption of imprinting caused by deletion of the H19 gene region in mice. Nature. 375, 34–39.

Leighton, P.A., Saam, J.R., Ingram, R.S., Stewart, C.L. and Tilghman, S.M. (1995b). An enhancer deletion affects both H19 and Igf2 expression. Genes & development. 9, 2079–2089.

Lescisin, K.R., Varmuza, S. and Rossant, J. (1988). Isolation and characterization of a novel trophoblast-specific cDNA in the mouse. Genes & development. 2, 1639–1646.

Li, H.Y., Chang, S.P., Yuan, C.C., Chao, H.T., Ng, H.T. and Sung, Y.J. (2003). Induction of p38 mitogen-activated protein kinase-mediated apoptosis is involved in outgrowth of trophoblast cells on endometrial epithelial cells in a model of human trophoblast-endometrial interactions. Biology of reproduction. 69, 1515–1524.

Liao, S., Vickers, M.H., Taylor, R.S., Jones, B., Fraser, M., McCowan, L.M., Baker, P.N. and Perry, J.K. (2017). Maternal serum IGF-1, IGFBP-1 and 3, and placental growth hormone at 20 weeks’ gestation in pregnancies complicated by preeclampsia. Pregnancy hypertension. 10, 149–154.

Lisboa, F.A., Warren, J., Sulkowski, G., Aparicio, M., David, G., Zudaire, E. and Dveksler, G.S. (2011). Pregnancy-specific glycoprotein 1 induces endothelial tubulogenesis through interaction with cell surface proteoglycans. Journal of Biological Chemistry. 286, 7577–7586.

Livak, K.J. and Schmittgen, T.D. (2001). Analysis of relative gene expression data using real-time quantitative PCR and the 2-ΔΔCT method. methods. 25, 402–408.

Lopez, M.F., Dikkes, P., Zurakowski, D.A.V.I.D. and Villa-Komaroff, L. (1996). Insulin-like growth factor II affects the appearance and glycogen content of glycogen cells in the murine placenta. Endocrinology. 137, 2100–2108.

López-Tello, J., Pérez-García, V., Khaira, J., Kusinski, L.C., Cooper, W.N., Andreani, A., Grant, I., de Liger, E.F., Lam, B.Y., Hemberger, M., et al. (2019). Fetal and trophoblast PI3K p110α have distinct roles in regulating resource supply to the growing fetus in mice. Elife. 8, e45282.

Mann, J.R. and Lovell-Badge, R.H. (1984). Inviability of parthenogenones is determined by pronuclei, not egg cytoplasm. Nature. 310, 66–67.

McGrath, J. and Solter, D. (1984). Completion of mouse embryogenesis requires both the maternal and paternal genomes. Cell. 37, 179–183.

McLellan, A.S., Fischer, B., Dveksler, G., Hori, T., Wynne, F., Ball, M., Okumura, K., Moore, T. and Zimmermann, W. (2005). Structure and evolution of the mouse pregnancy-specific glycoprotein (Psg) gene locus. BMC genomics. 6, 1–17.

Moore, T. and Haig, D. (1991). Genomic imprinting in mammalian development: a parental tug-of-war. Trends in genetics. 7, 45–49.

Morgan, D.O., Edman, J.C., Standring, D.N., Fried, V.A., Smith, M.C., Roth, R.A. and Rutter, W.J. (1987). Insulin-like growth factor II receptor as a multifunctional binding protein. Nature. 329, 301–307.

Mudgett, J.S., Ding, J., Guh-Siesel, L., Chartrain, N.A., Yang, L., Gopal, S. and Shen, M.M. (2000). Essential role for p38α mitogen-activated protein kinase in placental angiogenesis. Proceedings of the National Academy of Sciences. 97, 10454–10459.

Musial, B., Fernandez-Twinn, D.S., Vaughan, O.R., Ozanne, S.E., Voshol, P., Sferruzzi-Perri, A.N. and Fowden, A.L. (2016). Proximity to delivery alters insulin sensitivity and glucose metabolism in pregnant mice. Diabetes. 65, 851–860.

Napso, T., Yong, H.E., Lopez-Tello, J. and Sferruzzi-Perri, A.N. (2018). The role of placental hormones in mediating maternal adaptations to support pregnancy and lactation. Frontiers in physiology. 9, 1091.

Napso, T., Zhao, X., Ibañez Lligoña, M., Sandovici, I., Kay, R., Gribble, F., Reimann, F., Meek, C., Hamilton, R.S. and Sferruzzi-Perri, A. (2020). Unbiased placental secretome characterization identifies candidates for pregnancy complications. bioRxiv. doi: 10.1101/2020.07.12.198366.

Ngala, R.A., Fondjo, L.A., Gmagna, P., Ghartey, F.N. and Awe, M.A. (2017). Placental peptides metabolism and maternal factors as predictors of risk of gestational diabetes in pregnant women. A case-control study. PloS one. 12, p.e0181613.

Nordin, M., Bergman, D., Halje, M., Engström, W. and Ward, A. (2014). Epigenetic regulation of the Igf2/H19 gene cluster. Cell proliferation. 47, 189–199.

Rakoczy, J., Padmanabhan, N., Krzak, A.M., Kieckbusch, J., Cindrova-Davies, T. and Watson, E.D. (2017). Dynamic expression of TET1, TET2, and TET3 dioxygenases in mouse and human placentas throughout gestation. Placenta. 59, 46–56.

Redline, R.W., Chernicky, C.L., Tan, H.Q., Ilan, J. and Ilan, J. (1993). Differential expression of insulin-like growth factor-II in specific regions of the late (post day 9.5) murine placenta. Molecular reproduction and development. 36, 121–129.

Ripoche, M.A., Kress, C., Poirier, F. and Dandolo, L. (1997). Deletion of the H19 transcription unit reveals the existence of a putative imprinting control element. Genes & development. 11, 1596–1604.

Rodgers, A. and Sferruzzi-Perri, A.N. (2021). Developmental programming of offspring adipose tissue biology and obesity risk. International Journal of Obesity. 1–23.

Rosenfeld, C.S. (2015). Sex-specific placental responses in fetal development. Endocrinology. 156, 3422–3434.

Salazar-Petres, E.R. and Sferruzzi-Perri, A.N. (2021). Pregnancy-induced changes in ß-cell function: what are the key players? The Journal of Physiology.

Sandovici, I., Georgopoulou, A., Hufnagel, A.S., Schiefer, S.N., Santos, F., Hoelle, K., Lam, B.Y., Yeo, G.S., Burling, K., López-Tello, J., et al. (2019). Fetus-derived IGF2 matches placental development to fetal demand. bioRxiv. http://doi.org/10.1101/520536

Sasaki, H., Ishihara, K. and Kato, R. (2000). Mechanisms of Igf2/H19 imprinting: DNA methylation, chromatin and long-distance gene regulation. The Journal of Biochemistry. 127, 711–715.

Schoenherr, C.J., Levorse, J.M. and Tilghman, S.M. (2003). CTCF maintains differential methylation at the Igf2/H19 locus. Nature genetics. 33, 66–69.

Schulz, R., Menheniott, T.R., Woodfine, K., Wood, A.J., Choi, J.D. and Oakey, R.J. (2006). Chromosome-wide identification of novel imprinted genes using microarrays and uniparental disomies. Nucleic acids research. 34, e88.

Sferruzzi-Perri, A.N. (2018). Regulating needs: Exploring the role of insulin-like growth factor-2 signalling in materno-fetal resource allocation. Placenta. 64, S16–S22.

Sferruzzi-Perri, A.N. and Camm, E.J. (2016). The programming power of the placenta. Frontiers in physiology. 7, 33.

Sferruzzi-Perri, A.N., Macpherson, A.M., Roberts, C.T. and Robertson, S.A. (2009). Csf2 null mutation alters placental gene expression and trophoblast glycogen cell and giant cell abundance in mice. Biology of reproduction. 81, 207–221.

Sferruzzi-Perri, A.N., Vaughan, O.R., Coan, P.M., Suciu, M.C., Darbyshire, R., Constancia, M., Burton, G.J. and Fowden, A.L. (2011). Placental-specific Igf2 deficiency alters developmental adaptations to undernutrition in mice. Endocrinology. 152, 3202–3212.

Sferruzzi-Perri, A.N., Vaughan, O.R., Forhead, A.J. and Fowden, A.L. (2013a). Hormonal and nutritional drivers of intrauterine growth. Current Opinion in Clinical Nutrition & Metabolic Care. 16, 298–309.

Sferruzzi-Perri, A.N., Vaughan, O.R., Haro, M., Cooper, W.N., Musial, B., Charalambous, M., Pestana, D., Ayyar, S., Ferguson-Smith, A.C., Burton, G.J., et al. (2013b). An obesogenic diet during mouse pregnancy modifies maternal nutrient partitioning and the fetal growth trajectory. The FASEB Journal. 27, 3928–3937.

Sferruzzi-Perri, A.N., López-Tello, J., Fowden, A.L. and Constancia, M. (2016). Maternal and fetal genomes interplay through phosphoinositol 3-kinase (PI3K)-p110α signaling to modify placental resource allocation. Proceedings of the National Academy of Sciences. 113, 11255–11260.

Sferruzzi-Perri, A.N., Sandovici, I., Constancia, M. and Fowden, A.L. (2017). Placental phenotype and the insulinlike growth factors: resource allocation to fetal growth. The Journal of physiology. 595, 5057–5093.

Sferruzzi-Perri, A.N., Lopez-Tello, J., Napso, T. and Yong, H.E. (2020). Exploring the causes and consequences of maternal metabolic maladaptations during pregnancy: Lessons from animal models. Placenta. 98, 43–51.

Sharma, N., Kubaczka, C., Kaiser, S., Nettersheim, D., Mughal, S.S., Riesenberg, S., Hölzel, M., Winterhager, E. and Schorle, H. (2016). Tpbpa-Cre-mediated deletion of TFAP2C leads to deregulation of Cdkn1a, Akt1 and the ERK pathway, causing placental growth arrest. Development. 143, 787–798.

Sharma, A., Lacko, L.A., Argueta, L.B., Glendinning, M.D. and Stuhlmann, H. (2019). miR-126 regulates glycogen trophoblast proliferation and DNA methylation in the murine placenta. Developmental biology. 449, 21–34.

Sibley, C.P., Coan, P.M., Ferguson-Smith, A.C., Dean, W., Hughes, J., Smith, P., Reik, W., Burton, G.J., Fowden, A.L. and Constancia, M. (2004). Placental-specific insulin-like growth factor 2 (Igf2) regulates the diffusional exchange characteristics of the mouse placenta. Proceedings of the National Academy of Sciences. 101, 8204–8208.

Siddle, K. (2011). Signalling by insulin and IGF receptors: supporting acts and new players. Journal of molecular endocrinology. 47, R1–R10.

Simmons, D.G., Fortier, A.L. and Cross, J.C. (2007). Diverse subtypes and developmental origins of trophoblast giant cells in the mouse placenta. Developmental biology. 304, 567–578.

Simmons, D.G., Rawn, S., Davies, A., Hughes, M. and Cross, J.C. (2008). Spatial and temporal expression of the 23 murine Prolactin/Placental Lactogen-related genes is not associated with their position in the locus. BMC genomics. 9, 1–20.

Snyder, S.K., Wessner, D.H., Wessells, J.L., Waterhouse, R.M., Wahl, L.M., Zimmermann, W. and Dveksler, G.S. (2001). Pregnancy-specific Glycoproteins Function as Immunomodulators by Inducing Secretion of IL-10, IL-6 and TGF-beta1 by Human Monocytes. American journal of reproductive immunology. 45, 205–216.

Srivastava, M., Hsieh, S., Grinberg, A., Williams-Simons, L., Huang, S.P. and Pfeifer, K. (2000). H19 and Igf2 monoallelic expression is regulated in two distinct ways by a shared cis acting regulatory region upstream of H19. Genes & development. 14, 1186–1195.

Su, R., Wang, C., Feng, H., Lin, L., Liu, X., Wei, Y. and Yang, H. (2016). Alteration in expression and methylation of IGF2/H19 in placenta and umbilical cord blood are associated with macrosomia exposed to intrauterine hyperglycemia. PloS one. 11, e0148399.

Surani, M.A., Barton, S.C. and Norris, M.L. (1984). Development of reconstituted mouse eggs suggests imprinting of the genome during gametogenesis. Nature. 308, 548–550.

Szabó, P.E., Tang, S.H.E., Rentsendorj, A., Pfeifer, G.P. and Mann, J.R. (2000). Maternal-specific footprints at putative CTCF sites in the H19 imprinting control region give evidence for insulator function. Current Biology. 10, 607–610.

Thorvaldsen, J.L., Duran, K.L. and Bartolomei, M.S. (1998). Deletion of the H19 differentially methylated domain results in loss of imprinted expression of H19 and Igf2. Genes & development. 12, 3693–3702.

Tremblay, K.D., Duran, K.L. and Bartolomei, M.S. (1997). A 5’2-kilobase-pair region of the imprinted mouse H19 gene exhibits exclusive paternal methylation throughout development. Molecular and cellular biology. 17, 4322–4329.

Tucci, V., Isles, A.R., Kelsey, G., Ferguson-Smith, A.C., Bartolomei, M.S., Benvenisty, N., Bourc’his, D., Charalambous, M., Dulac, C., Feil, R., et al. (2019). Genomic imprinting and physiological processes in mammals. Cell. 176, 952–965.

Tunster, S.J., Jensen, A.B. and John, R.M. (2013). Imprinted genes in mouse placental development and the regulation of fetal energy stores. Reproduction. 145, R117–R137.

Tunster, S.J., Creeth, H.D.J. and John, R.M. (2015). The imprinted Phlda2 gene modulates a major endocrine compartment of the placenta to regulate placental demands for maternal resources. Developmental biology. 409, 251–260.

Tunster, S.J., McNamara, G.I., Creeth, H.D.J. and John, R.M. (2016). Increased dosage of the imprinted Ascl2 gene restrains two key endocrine lineages of the mouse Placenta. Developmental biology. 418, 55–65.

Tunster, S.J., Boqué-Sastre, R., McNamara, G.I., Hunter, S.M., Creeth, H.D. and John, R.M. (2018). Peg3 deficiency results in sexually dimorphic losses and gains in the normal repertoire of placental hormones. Frontiers in cell and developmental biology. 6, 123.

Tunster, S.J., Watson, E.D., Fowden, A.L. and Burton, G.J. (2020). Placental glycogen stores and fetal growth: insights from genetic mouse models. Reproduction. 159, R213–R235.

Verona, R.I., Mann, M.R. and Bartolomei, M.S. (2003). Genomic imprinting: intricacies of epigenetic regulation in clusters. Annual review of cell and developmental biology. 19, 237–259.

Wada, T., Hori, S., Sugiyama, M., Fujisawa, E., Nakano, T., Tsuneki, H., Nagira, K., Saito, S. and Sasaoka, T. (2010). Progesterone inhibits glucose uptake by affecting diverse steps of insulin signaling in 3T3-L1 adipocytes. American Journal of Physiology-Endocrinology and Metabolism. 298, 881–888.

Wislocki, G.B. and Bennett, H.S. (1943). The histology and cytology of the human and monkey placenta, with special reference to the trophoblast. American Journal of Anatomy. 73, 335–449.

Wu, J.A., Johnson, B.L., Chen, Y., Ha, C.T. and Dveksler, G.S. (2008). Murine pregnancy-specific glycoprotein 23 induces the proangiogenic factors transforming-growth factor beta 1 and vascular endothelial growth factor a in cell types involved in vascular remodeling in pregnancy. Biology of reproduction. 79, 1054–1061.

